# Metabolic rescue ameliorates mitochondrial encephalo-cardiomyopathy in murine and human iPSC models of Leigh syndrome

**DOI:** 10.1101/2022.01.04.475008

**Authors:** Jin-Young Yoon, Nastaran Daneshgar, Yi Chu, Biyi Chen, Marco Hefti, Kaikobad Irani, Long-Sheng Song, Charles Brenner, E. Dale Abel, Barry London, Dao-Fu Dai

**Author notes:** Corresponding author: Dao-Fu Dai, MD, PhD Department of Pathology, University of Iowa Carver College of Medicine 25 S. Grand Ave, Iowa City, IA 52242. Room MRC106C. Co-first authors (Jin-Young Yoon, Nastaran Daneshgar). **Conflict of interests:** The authors have declared that no conflict of interest exists.

## Abstract

Mice with deletion of complex I subunit Ndufs4 develop mitochondrial encephalomyopathy resembling Leigh syndrome (LS). We report that LS mice also develop severe cardiac bradyarrhythmia and diastolic dysfunction. Human induced pluripotent stem cell-derived cardiomyocytes (iPS-CMs) with Ndufs4 deletion recapitulate LS cardiomyopathy. Mechanistically, we demonstrate a direct link between complex I deficiency, decreased intracellular NAD^+^/ NADH and bradyarrhythmia, mediated by hyperacetylation of the cardiac sodium channel Na_V_1.5, particularly at K1479 site. Neuronal apoptosis in the cerebellar and midbrain regions in LS mice was associated with hyperacetylation of p53 and activation of microglia. Targeted metabolomics revealed increases in several amino acids and citric acid cycle intermediates, likely due to impairment of NAD^+^-dependent dehydrogenases, and a substantial decrease in reduced Glutathione (GSH). Metabolic rescue by nicotinamide riboside (NR) supplementation increased intracellular NAD^+^/ NADH, restored metabolic derangement, reversed protein hyperacetylation through NAD^+^-dependent Sirtuin deacetylase, and ameliorated cardiomyopathic phenotypes, concomitant with improvement of Na_V_1.5 current and SERCA2a function measured by Ca2^+^- transients. NR also attenuated neuronal apoptosis and microglial activation in the LS brain and human iPS-derived neurons with Ndufs4 deletion. Our study reveals direct mechanistic explanations of the observed cardiac bradyarrhythmia, diastolic dysfunction and neuronal apoptosis in mouse and human iPSC models of LS.

## INTRODUCTION

Leigh syndrome (LS) is a severe mitochondrial disorder that manifests as psychomotor regression early in life (1), and treatment options are very limited, as is the case for many mitochondrial disorders. The most common causes are mutations in components of mitochondrial complex I, NADH dehydrogenase, which consists of 45 subunits, one of which is NADH dehydrogenase [ubiquinone] iron-sulfur protein 4 (Ndufs4). Mice with a homozygous germline deletion of exon 2 of the encoding gene (*Ndufs4^-/-^*) exhibit phenotypes resembling the severe encephalomyopathy in LS patients (2). These include features of growth retardation, lethargy, loss of motor skills, ataxia, hypothermia, slowed breathing and apnea. The latter has been shown to contribute to early death (∼50-60 days after birth) in this mouse model.

Among LS patients, approximately 18-21% have cardiac involvement and this is associated with a worse prognosis. Cardiac abnormalities in LS may include cardiomyopathy, pericardial effusion, and conduction abnormalities. Hypertrophic cardiomyopathy has been reported as the most common abnormality in these patients *(3-4)*. In Ndufs4^-/-^ mouse models, while the encephalomyopathy is unequivocal, cardiac involvement remains debatable. In two independent studies describing mice with cardiac-specific Ndufs4 deletion, one showed hypertrophic cardiomyopathy (5), while the other reported normal cardiac function (6).

In the present study, we sought to elucidate the metabolic derangement and underlying mechanisms of cardio-encephalomyopathy in LS using germline Ndufs4^-/-^ mice *(henceforth referred to as LS mice*). We found that these mice develop severe bradyarrhythmia and diastolic dysfunction due to hyperacetylation of the cardiac sodium channel (Na_V_1.5) and the calcium handling protein sarco/endoplasmic reticulum Ca^2+^-ATPase 2a (SERCA2a), respectively. We also discovered that supplementation with nicotinamide riboside (NR) (7) increased intracellular NAD^+^, reversed protein hyperacetylation, ameliorated metabolic derangement, restored sinus rhythm, normalized diastolic function, ameliorated the decreases in Na_V_1.5 current and restored SERCA2 function in LS mice and human iPS-CMs with homozygous Ndufs4 deletion. We further demonstrate that NR attenuates acetyl-p53-mediated neuronal apoptosis in LS mouse brain (cerebellum and midbrain) and iPS-derived mixed neurons with the Ndufs4 deletion. These data provide a mechanistic explanation for the pathophysiology of LS cardio-encephalomyopathy and its underlying metabolic derangement.

## RESULTS

### Metabolic derangement in LS is partially restored by Nicotinamide Riboside

We performed multiple targeted metabolomics analyses to investigate changes of several metabolites in the LS heart and brainstem/cerebellum. We focused on brainstem and cerebellum since these regions have the most prominent brain pathologies and can be quickly dissected and flash-frozen to preserve tissue metabolites. The metabolomic dataset is presented in Supplemental Table 1 (1^st^, WT vs. LS hearts), Supplemental Table 2 (2^nd^, WT vs. LS vs. NR-treated LS hearts), Supplemental Table 3 (2^nd^, WT vs. LS vs. NR-treated LS brainstem and cerebella) and is summarized in Figure 1a-b. The most prominent findings were significant increases in many amino acids (ranged from +36% to >3-fold increase in both hearts and brains), urea and metabolic derivatives-keto-isocaproate and hydroxy-β-methyl-of branched-chain amino acid (BCAA) aminotransferase, including α-keto-isocaproate and β hydroxy-β-methyl1 butyric acid (both from leucine), α-keto-isovalerate (KIV, from valine) and α-keto-β-methylvaleric acid (KMV, from isoleucine). These α-keto-derivatives of BCAA usually undergo oxidation by branched-chain α-keto acid dehydrogenases (BCKD), key enzymes requiring NAD^+^ as a cofactor. BCKD enzymes were likely impaired because of decreased NAD^+^/NADH in LS hearts and brains. There were accumulation of some TCA cycle intermediates, including citrate, aconitate, and isocitrate, which is likely related to impaired activity of the NAD^+^-dependent enzyme isocitrate dehydrogenase. Arachidonate, a critical signaling molecule and a precursor of various eicosanoids (such as prostaglandins, etc.), was increased. There were decreases in niacin (a precursor of NAD^+^), tyrosine and mannose (only in the heart). NR supplementation (500mg/Kg/day, i.p.) significantly reversed and restored many of these metabolomics changes associated with Ndufs4 deficiency in both cerebellum/brainstem and heart (Figure 1a-b). There were no significant and consistent changes in glycolytic, pentose phosphate pathways, purine, or pyrimidine metabolites.

**Fig. 1.**
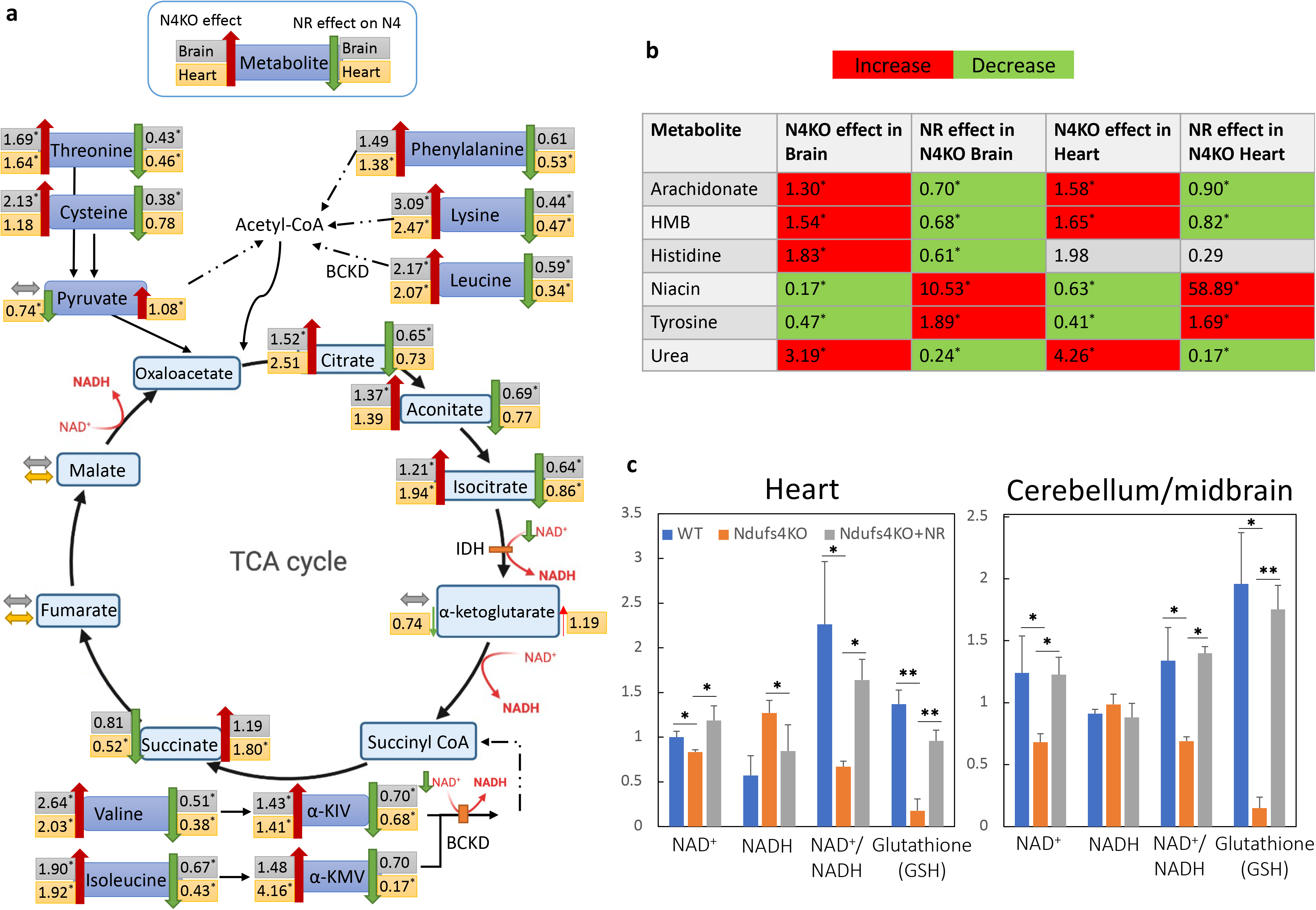
Metabolomic derangement and the effect of Nicotinamide Riboside. (**a**) Ratios of several metabolites in brain and heart of Ndufs4KO/WT (N4KO effect, left, n=4) and NR-treated Ndufs4KO/untreated Ndufs4KO (NR effect, right, n=3). IDH: Isocitrate dehydrogenase, BCKD: branched-chain α-keto acid dehydrogenase, α-KIV: ketoisovalerate, α-KMV: α-keto-β-methylvaleric acid. (**b**) Relative abundance of significantly changed metabolites in the indicated groups. HMB: beta-Hydroxy-beta-Methylbutyric-acid. (**c**) Representative metabolites data measured by Liquid chromatography /mass spectrometry in LS heart and cerebellum/midbrain. Data are mean ± s.e.m. of biologically independent samples. Statistical significance was determined by Student’s t-test; n=3-4 each group. (* p<0.05, **p<0.01).

Using liquid chromatography followed by mass spectrometry analysis of these samples, we demonstrated that Ndufs4 deletion (which results in the deficiency of NADH Dehydrogenase) led to decreased NAD^+^/NADH (Figure 1c). In addition, there was substantial depletion of the reduced form of glutathione (GSH), a major intracellular antioxidant system, in LS hearts and cerebellum. NR supplementation significantly rescued NAD^+^/ NADH and restored GSH levels in LS hearts and brains (Figure 1c), thereby ameliorated many metabolic defects due to NAD^+^ deficiency and redox stress (Figure 1a-b).

### Ndufs4^-/-^ mice (LS mice) were runted and had cardiomyopathy

Echocardiography of conscious mice showed that left ventricle (LV) mass and ejection fraction were normal in LS mice (Figure 2a-b), but tissue Doppler imaging showed that diastolic function was impaired (Figure 2c), as evidenced by decreased ratio of relaxation velocity of mitral annulus during early diastole (E’) to late diastole (A’). In addition, these mice had severe bradycardia, with a median resting heart rate (HR) of ∼400 bpm (Figure 2d), reflecting a ∼35% decrease from the baseline physiological HR of >600 bpm. Both diastolic dysfunction and bradyarrhythmia were significantly ameliorated by supplementation with NR (500mg/Kg/day, i.p.) (Figure 2c-d). LS mice were runted, with their average body weight ∼60% that of wild type (WT) littermates, and NR supplementation did not alter this (Supplemental Figure 1a). Heart weight and heart weight normalized to tibia length did not differ significantly from that in WT mice, and this was not affected by NR supplementation (Supplemental Figure 1b-c). Pathological analysis assessing ventricular size and fibrosis did not distinguish between hearts from WT mice and their LS counterparts with or without NR treatment (Supplemental Figure 1d-g).

**Fig. 2.**
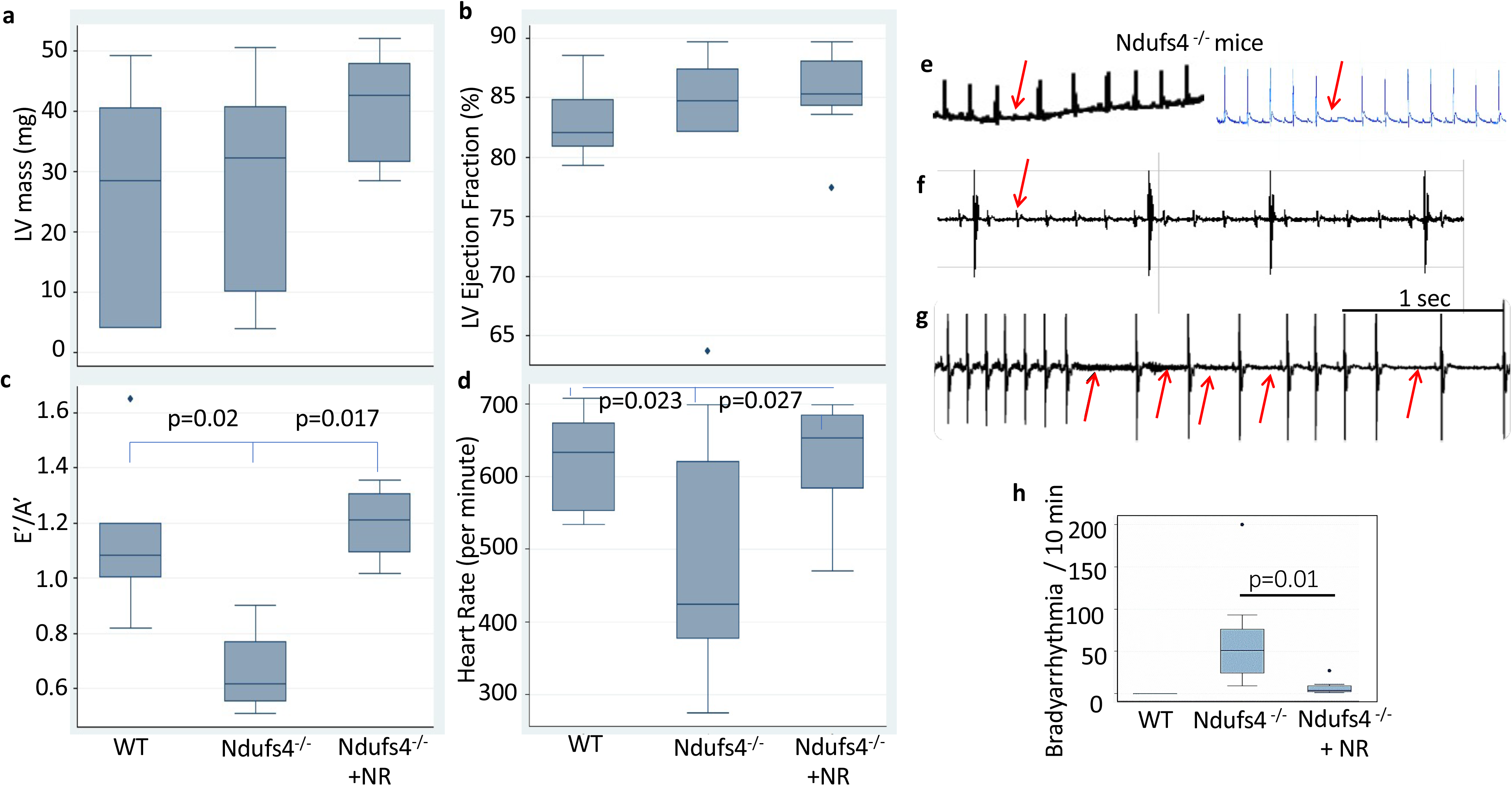
Diastolic dysfunction and bradyarrhythmia in Ndufs4 deficient mice is reversed by treatment with nicotinamide riboside (NR). Echocardiographic measurements of **(a)** left ventricle mass (LV), **(b)** LV ejection fraction, **(c)** E’/A’ as determined by tissue Doppler imaging (impaired diastolic function defined as È/À<1), **(d)** Heart rate (n=8-10). **(e-g)** Representative ECG of Ndufs4^-/-^ mice, showing **(e)** episodic second-degree atrioventricular (AV) block, **(f)** third-degree AV block, and **(g)** sinus arrhythmia. **(h)** Quantitation of arrhythmic events over a 10-minute period in wild type, Ndufs4^-/-^ (LS) mice, with or without NR treatment.

Electrocardiograms (ECGs) from conscious immobilized LS mice demonstrate a spectrum of bradyarrhythmia. These include occasional second-degree atrioventricular (AV) block (red arrow, Figure 2e), rare high-grade AV block (Figure 2f), and frequent sinus node dysfunction (Figure 2g). Bradyarrhythmia were frequently recorded in LS mice (Figure 2h). Supplementation with NR substantially reduced arrhythmic events in LS mice (Figure 2h).

### Hyperacetylation of Na_V_1.5 promote bradyarrhythmia in LS mice

To exclude autonomic dysfunction or metabolic acidosis as potential contributors to the bradyarrhythmia observed in LS mice, we dissected sinoatrial nodal (SAN) tissue, loaded it with the Rhod-2 calcium indicator dye *ex vivo*, and assessed calcium transients by confocal microscopy. In LS SAN tissue, there were multiple long pauses in spontaneous [Ca^2+^] transients, suggesting SA nodal pauses (Figure 3b). NR supplementation significantly reduced the bradycardic events and restored sinus rhythm (Figure 3c-d) to frequencies and amplitudes similar to those recorded from WT SAN samples (Figure 3a). Fibrosis assessed by Trichome stain was not different in WT SAN, LS SAN or LS SAN treated with NR (Figure 3e-g).

**Fig. 3.**
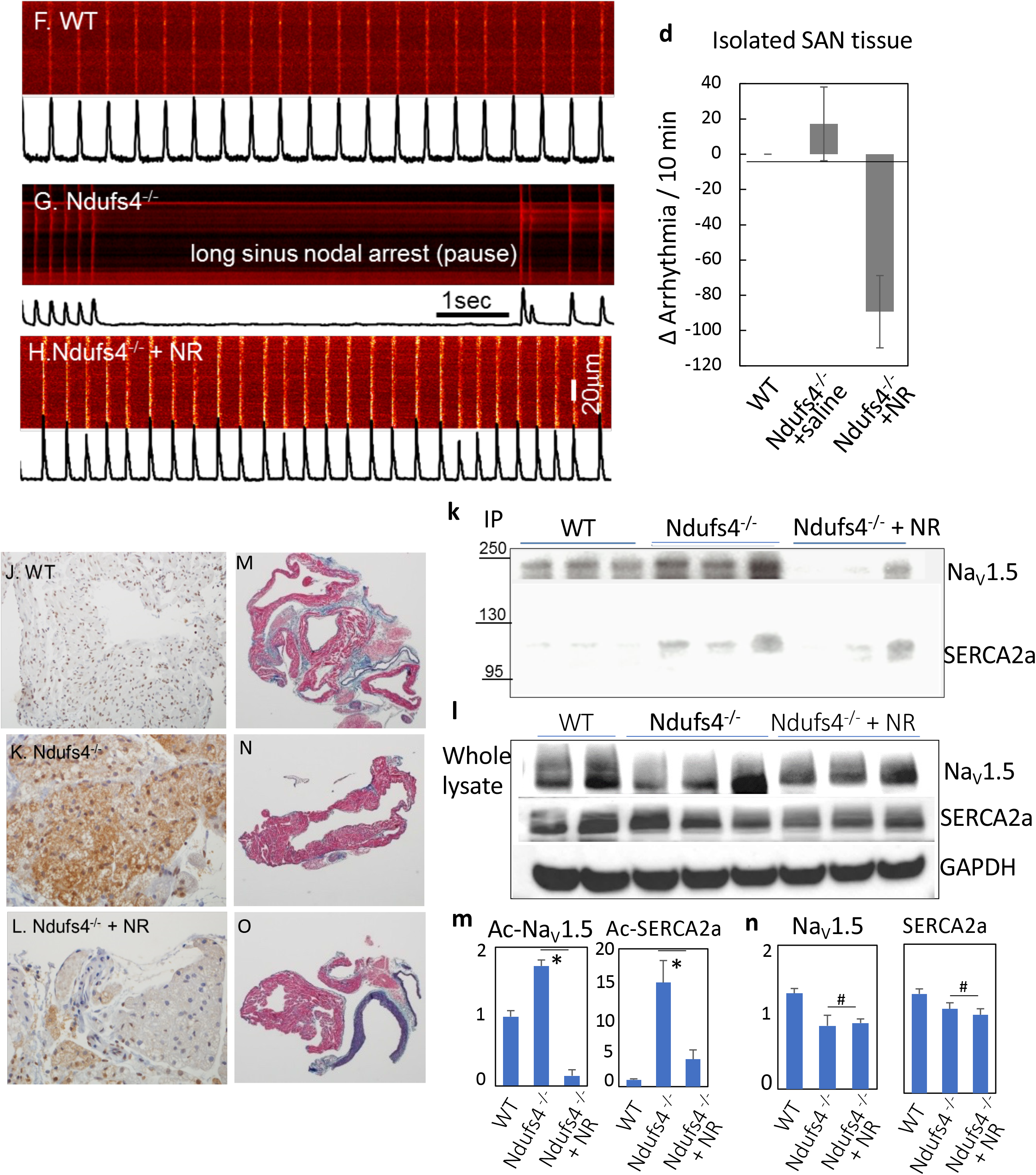
Characteristics and mechanism of bradyarrhythmia in Ndufs4*^-/-^* Leigh Syndrome (LS) mice. (**a-j**) Analyses of sinoatrial node (SAN) tissue dissected from the hearts of mice from three designated groups. Confocal imaging of Ca^2+^-transients in SANs tissue from (**a**) WT, (**b**) Ndufs4^-/-^ and (**c**) Ndufs4^-/-^mice treated with nicotinamide riboside (NR). (**d**) Quantitation of changes in arrhythmic events in SANs over a 10-minute period (post-pretreatment). Trichrome staining of SANs in (**e**) WT tissue, (**f**) Ndufs4^-/-^ tissue, and (**g**) Ndufs4^-/-^ tissue from NR-treated mice. Immunohistochemical staining of SANs with specific antibody against acetylated K1479-Na_V_1.5 in (**h**) WT tissue, (**i**) Ndufs4^-/-^ tissue, and (**j**) NR-treated Ndufs4^-/-^ tissue. (**k**) Western blot of mouse heart proteins from specified treatment groups following immunoprecipitation with total acetyl-lysine antibodies and probing with Na_V_1.5 or SERCA2a antibodies. (**l**) Total Na_V_1.5 and SERCA2a proteins in mouse hearts of the indicated experimental groups. (**m-n**) Quantification of acetylated and total proteins in the indicated groups (* p<0.05, # p not significant). Data are mean ± s.e.m. of biologically independent samples. Statistical significance was determined by Student’s t-test; n=3-4 for immunoblots.

To address whether cell-autonomous mechanism is sufficient to cause arrhythmia, we crossed Ndufs4^flox/flox^ mice with HCN4-Cre mice to generate conduction tissue-specific deletion of Ndufs4. Approximately 2-3 months after tamoxifen-induced deletion of Ndufs4, 87.5% of the HCN4-Ndufs4^-/-^ mice developed various forms of arrhythmia, including atrial or ventricular premature contractions, sinus bradyarrhythmia and frequent sinus pauses resembling those found in germline Ndufs4^-/-^ LS mice (Supplemental Figure 2a-c). This finding supports the hypothesis that Ndufs4 deficiency in the cardiac conduction system itself is sufficient to cause arrhythmia in a cell-autonomous manner, although the metabolic defect of Ndufs4^-/-^ in neurons, cardiomyocytes, and other cell types could also contribute to arrhythmia by impacting the neurohormonal or paracrine regulation of cardiac rhythm.

Ndufs4 deletion (which results in the deficiency of NADH Dehydrogenase) led to reduced NAD^+^/NADH (Figure 1c), a change that can impair the function of NAD^+^-dependent enzymes, including Sirt1(8). To identify potential mechanisms responsible for the arrhythmia and diastolic dysfunction in LS mice, we examined the cardiac sodium channel Na_V_1.5 and SERCA2a, respectively. Na_V_1.5 governs the initiation and propagation of the cardiac action potential, whereas SERCA2a is the key protein regulating the decay of calcium transients by reuptake into sarcoplasmic reticulum. Both proteins are reported to be modulated by Sirt1^9,10^. We have previously shown that acetylation of the K1479 residue in Na_V_1.5 impairs the trafficking of this channel protein, causing a decrease in the inward depolarizing Na^+^ current (I_Na_) conducted by this channel. To examine the role of acetylation in LS hearts, we performed immunohistochemistry using an antibody specific for acetylation of Na_V_1.5 at K1479 modification and showed that K1479 in Na_V_1.5 was significantly hyperacetylated in both the SAN and LV from LS hearts (Figure 3i, Supplemental Figure 3a-b). NR supplementation dramatically reversed this acetylation in both SAN and LV in LS mouse hearts (Figure 3j, Supplemental Figure 3c), concomitant with the observed amelioration of arrhythmia and restoration of the normal sinus rhythm (Figure 3a-c). For quantitative analysis, we performed immunoprecipitation with acetyl-lysine antibody and demonstrated that the acetylated forms of Na_V_1.5 and SERCA2a were substantially increased in LS hearts. The acetylated Na_V_1.5 and SERCA2a were significantly attenuated by NR supplementation (Figure 3k, m), while the total levels of both the Na_V_1.5 and SERCA2a proteins did not differ significantly among any groups (Figure 3l, n).

### Acetylation of K1479 of Na_V_1.5 in mitochondrial complex I -deficient cells decreases the inward depolarizing Na^+^ current

To further investigate the role of K1479 acetylation in Na_V_1.5 in the context of mitochondrial complex I deficiency, we used HEK293 cells heterologous expression system. Since the endogenous Na_V_1.5 expression in HEK293 is negligible, we can investigate Na_V_1.5 by introducing exogenous wild-type or K1479 mutant (Figure 4a). We generated Ndufs4 and Ndufs2 knock-out (KO) HEK 293 cells using the CRISPR/Cas9 method, followed by transfection with WT or K1479 mutant Na_V_1.5, as confirmed by Western blots (Figure 4a). Notably, the deletion of Ndufs2 also resulted in a substantial decrease in Ndufs4 expression. Given that mitochondrial complex I is a large respiratory complex and Ndufs2 is located adjacent to the junction of the inner membrane and matrix domain, we proposed that its loss destabilized the matrix domain leading to degradation of the Ndufs4 subunit. As expected, in light of the loss of both subunits in the Ndufs2 KO cells, the phenotype was more severe than that of the Ndufs4 KO cells. Analysis of the conditioned media from these two cell lines and control cells, revealed that by 24 hours, the Ndufs2 KO cells developed a significantly more acidic environment (pH ∼7.2) than WT cells (pH ∼7.7, p=0.023), while the media from the Ndufs4 KO cells did not become significantly acidic until 48 hours (pH of ∼6.9) compared to WT cells (pH ∼7.3, p=0.01) and remained less acidic than Ndufs2 KO media (pH ∼6.5, p=0.006, Figure 4b). Immunoblotting confirmed elevated protein acetylation of total protein from Ndufs2KO cell lysate (Supplemental Figure 3d).

**Fig. 4.**
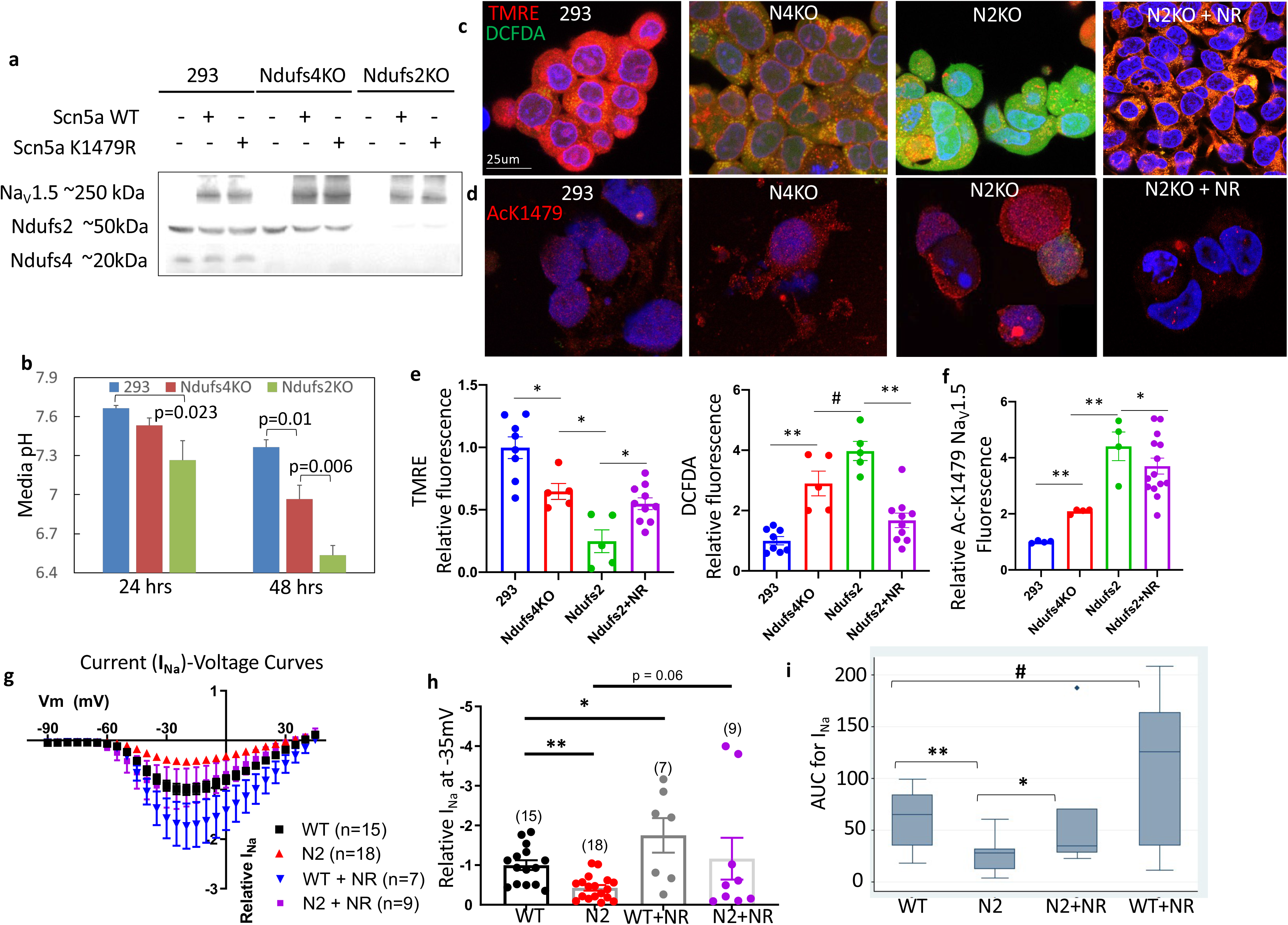
Acetylation of K-1479-Na_V_1.5 in mitochondrial complex I-deficient cells decrease sodium current. (**a**) Western blot for Na_V_1.5 protein in 293 cells lacking Ndufs4 (Ndufs4 KO) or Ndufs2 (Ndufs2 KO) which were transfected with plasmid carrying WT or mutant Scn5a gene (to produce WT or acetylation-site mutant Na_V_1.5 proteins). (**b**) pH of conditioned-media from 293, Ndufs4 KO, and Ndufs2 KO cells at 24 or 48 hours after replating. (**c**) Live staining of 293 (WT), Ndufs4 KO, and Ndufs2 KO cells with markers of mitochondrial membrane potential (TMRE) and total cellular ROS (DCFDA). (**d**) Representative fluorescence images of acetylation of Na_V_1.5 residue K1479. (**e**) Quantitation of results in panel *c*. (**f**) quantitation of results in *d*. (**g**) Patch clamp results of I_Na_ for WT and Ndufs2 KO cells (N2) with and without NR treatment. Quantitation of relative I_Na_ density at -35 mV (**h**) and AUC (**i**) for the indicated groups in panel *g*. n=7-18.

Live-cell staining with a marker of mitochondrial respiratory activity, tetramethylrhodamine, ethyl ester (TMRE), and a marker of reactive oxygen species (ROS), 2’,7’-dichlorodihydrofluorescein diacetate (DCFDA), showed reduced TMRE in Ndufs4 KO cells, indicating that mitochondrial membrane potential was decreased. Conversely, DCFDA fluorescence was increased, suggesting that total cellular ROS levels were higher (Figure 4c, e). Ndufs2 KO cells displayed a much greater loss of mitochondrial membrane potential and a greater increase in cellular ROS than the Ndufs4KO cells, and this was significantly ameliorated upon NR supplementation (Figure 4c, e). Immunofluorescence staining with an antibody specific for acetylated K1479 (Ac-K1479) in Na_V_1.5 revealed a ∼2-fold increase in acetylation in Ndufs4 KO cells and a ∼4-fold increase in Ndufs2 KO cells, which was significantly decreased after NR supplementation (Figure 4d, f). To determine the functional relevance of Na_V_1.5 hyperacetylation on this residue, we performed a patch-clamp experiment on the WT and Ndufs2 KO cells. The inward depolarizing Na^+^ current (I_Na_) was significantly lower in the Ndufs2 KO cells (Figure 4g-i), as shown by the I_Na_ at -35mV and the area under the current-voltage curves (AUC). Consistent with the contribution of Na_V_1.5 hyperacetylation (Figure 4d, f), pre-treatment with NR for 24hrs significantly restored I_Na_ in Ndufs2 KO cells and potentiated the I_Na_ of WT control cells (Figure 4g-i).

To further investigate the importance of K1479-Na_V_1.5 acetylation status on the Na^+^-current phenotype of Ndufs2KO cells, we transfected HEK293 cells with a lysine to glutamine mutant construct, K1479Q-Na_V_1.5, that mimics constitutively acetylated K1479 Na_V_1.5, which we previously showed to have decreased I_Na_ (9). Interestingly, when the K1479Q-Na_V_1.5 was expressed in Ndufs2 knock-out HEK293 cells, no further decrease in I_Na_ was observed compared with wild-type Na_V_1.5 in these Ndufs2KO cells (Supplemental Figure 4a-b).

These data indicate that acetylated K1479 Na_V_1.5 in these Ndufs2KO cells decreased I_Na_ to a similar extent as acetylation mimics K1479Q Na_V_1.5, suggesting a similar mechanism for both. While NR supplementation increased I_Na_ conducted by wild-type Na_V_1.5 (Figure 4h), the acetylation mimic K1479Q-Na_V_1.5 did not respond to NR in Ndufs2 KO cells (Supplemental Figure 4a-b). Since the K1479Q-Na_V_1.5 that mimics the persistent acetylated state led to decreased I_Na_ and did not respond to NR supplementation, these results support the hypothesis that acetylation of Na_V_1.5 at K1479 contributes to decreased I_Na_ induced by Ndufs2 deletion and that deacetylation by NR enhances I_Na_. To investigate whether the enhanced Na^+^ current phenotype by NR is mediated through Sirt1, we treated wild-type HEK293 cells expressing Na_V_1.5 with the Sirt1 inhibitor, Ex-527^9,11^. Our results showed that Ex-527 blocked the NR-mediated enhancement of I_Na_ (Supplemental Figure 4 c-d), suggesting that the NR effect is mediated, at least in part, by the NAD^+^-dependent enzyme Sirt1.

### NR ameliorates LS mice activity, neuronal apoptosis and microgliosis

Measurements performed in metabolic chambers showed that the LS mice had impaired motor function, characterized by cerebellar ataxia, and decreased locomotor activity (Figure 5a). Energy expenditure (heat production) and respiratory exchange ratio (RER) were lower in LS vs. WT mice (Figure 5b-c). These changes in energy expenditure were not significantly affected by NR. However, NR supplementation significantly ameliorated the ataxia (see Expanded View movies) and improved daily locomotor activity (Figure 5a).

**Fig. 5.**
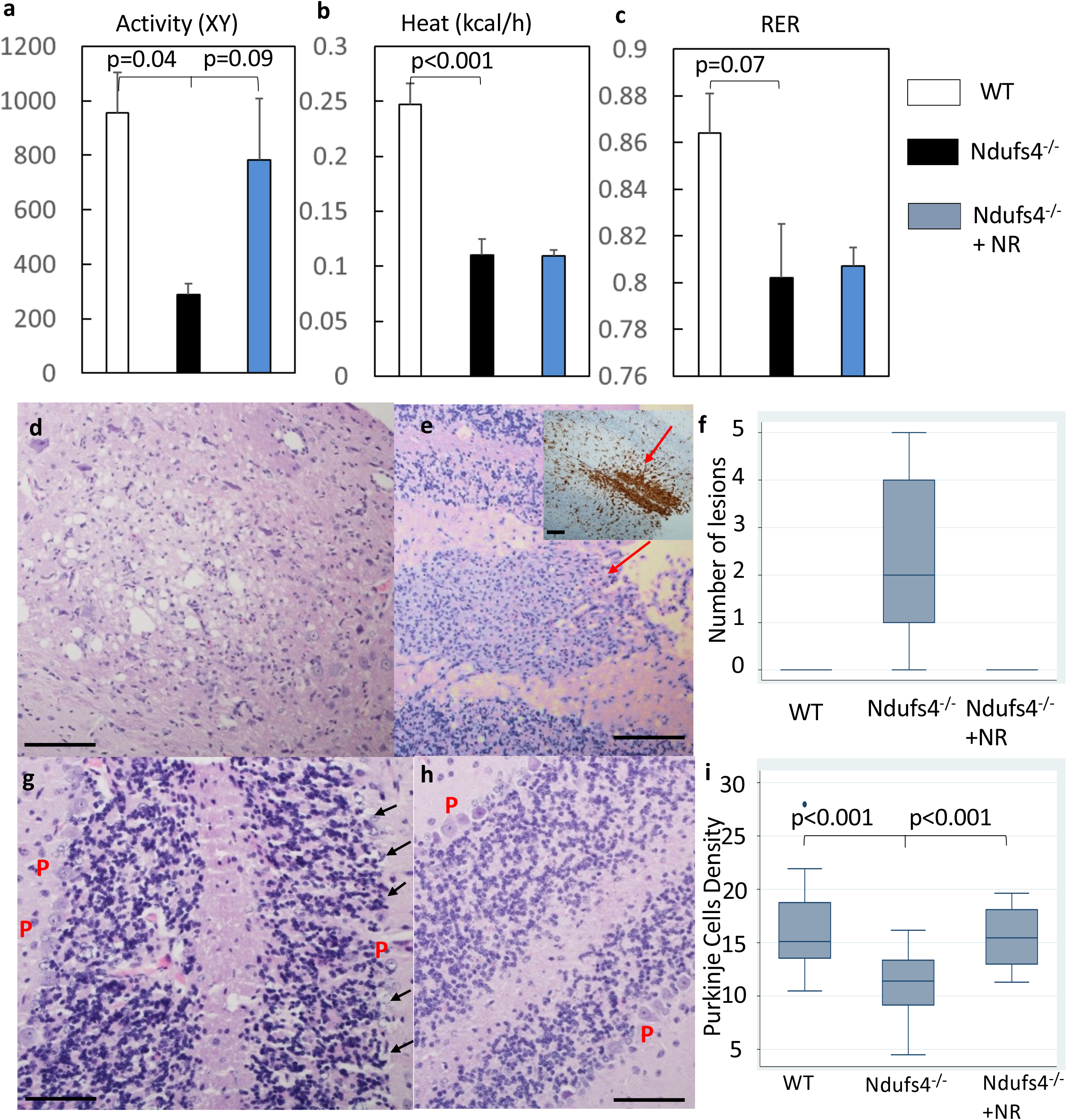
Neuronal apoptosis in cerebellum and cerebral cortex of LS mice. (**j-m**) Cerebellum and brainstem stained for activated caspase-3 per 10 high power fields (40x) (HPFs), for (**j**) WT, (**k**) Ndufs4^-/-^and (**l**) NR-treated Ndufs4^-/-^mice, (**m**) Quantitation of caspase-3 positive cells. (**n-q**) Cerebral cortex stained for activated caspase 3, and quantitation in mice of the same genotypes as in *j-m*. Inset in *o* is higher magnification image. (**r-u**) Iba-1 expression in the dorsal midbrain (**s**, arrow) and cerebellar cortex (**s**, arrowhead in the inset) of (**r**) WT, (**s**) Ndufs4^-/-^and (**t**) NR-treated Ndufs4^-/-^mice, (**u**) Quantitation of Iba-1 immunostaining; n=4-6; Kruskal Wallis test.+NR

**Fig. 5.**
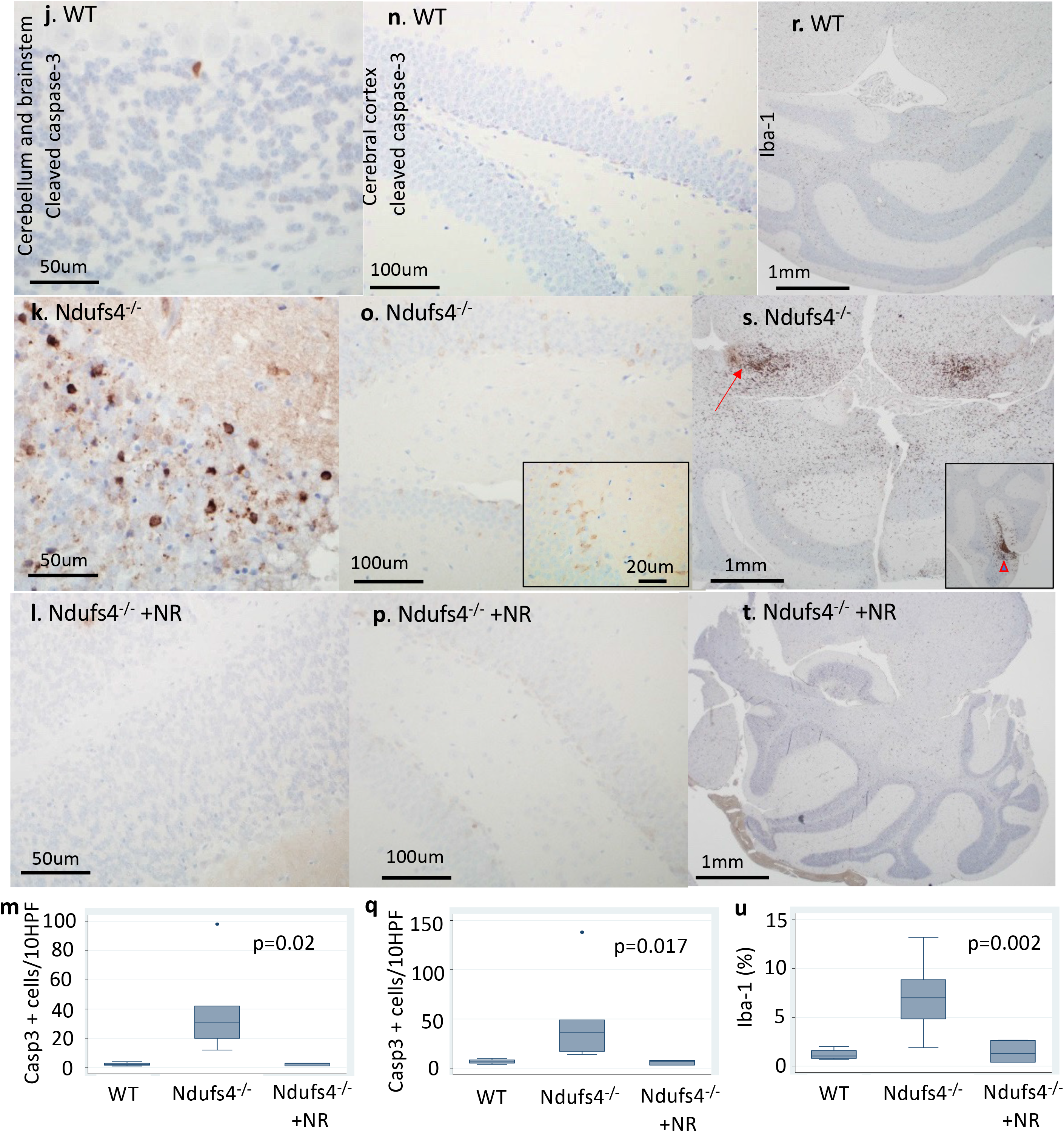
Metabolic Derangement and Neuropathology in LS mice. (**a-c**) Measurement of (**a**) mouse activity, (**b**) heat produced, and (**c**) respiratory exchange ratio (RER) using metabolic chambers; n=5-6. (**d-g**). Neuropathology of Ndufs4^-/-^ brain as detected by H&E staining of: (**d**) brainstem region, showing vacuolization of neuropils (spongiosis); (**e**) cerebellum, showing microglial and vascular proliferation (red arrow) (Iba-1 expression in the inset); (**f**) quantification of lesions described in *d* and *e*; (**g**) cerebellar cortex, showing Purkinje (P) cells (arrow). (**h**) Neuropathology of Ndufs4^-/-^ cerebellum following treatment with NR, as detected by H&E staining showing Purkinje cell preservation. (**i**) Quantification of Purkinje cell density in all three treatment groups. For all panels shown, Scale bar: 100 μm, N=4-5.

Neuropathological examination of the LS brain demonstrated spongiosis, defined as edema of neuropils (Figure 5d), and proliferation of microglia and vascular cells (Figure 5e, red arrow), particularly in the cerebellum and the brainstem region. Semiquantitative analysis of hematoxylin and eosin (H&E)-stained LS brain sections by a neuropathologist blinded to treatment group revealed increased foci of these lesions in the LS mice but none in NR-treated LS counterparts (Figure 5f). Quantitative analysis of Purkinje cell density in the cerebellar cortex showed a significant reduction of Purkinje cells in the LS mice, indicating that neurons were lost (Figure 5g, i), and this was attenuated by NR treatment (Figure. 5h-i).

Immunohistochemistry for cleaved-caspase-3 revealed prominent apoptosis among cells of the Purkinje layer and the underlying granular neurons in the cerebellum of untreated LS mice (Figure 5k, m). Apoptotic cells were scattered throughout the cerebral cortex in LS mice (Figure 5o, q). Apoptosis was associated with prominent activation of microglia, as the number of Iba-1 positive microglia in the dorsal midbrain (Figure 5s) and cerebellar cortex (Figure 5s inset) were significantly higher in the LS mice. Treatment with NR significantly mitigated neuronal apoptosis in both the cerebral cortex and dorsal midbrain (Figure 5l-m, 5p-q), and the associated activation of microglia in the LS brain (Figure 5t-u). Consistent with neuroprotective and cardioprotective effects, NR significantly extended the lifespan of LS mice (Supplemental Figure 5). Interestingly, although Ndufs4^-/-^ promoted apoptosis in the susceptible neurons of the cerebellar and brainstem regions, it did not induce apoptosis in cardiac left ventricular tissue (Supplemental Figure 6).

### Deletion of Ndufs4 decreased Na_V_1.5 current in human iPS-derived cardiomyocytes

To increase the translational relevance of our findings in mice and heterologous cells (HEK293), we deleted the Ndufs4 subunit using the CRISPR/Cas9 method in human iPS cells generated from a healthy Caucasian male (ATCC-1026). We confirmed that deletion of Ndufs4 was successful by Western blotting (Figure 6a) and immunostaining of the iPS-Cardiomyocytes (iPS-CMs) (Figure 6b). We performed live-cell staining of isogenic control and Ndufs4 KO iPS-CMs at ∼30 days post-differentiation. The presence of Ndufs4 staining in iPS-CM at this stage is confirmed by colocalization of Troponin staining in control iPS-CM (Supplemental Figure 7). Ndufs4 KO cells had less TMRE staining (loss of red) and more DCFDA staining (green) than control iPS-CMs, and this was mitigated by pre-treatment with NR (1mM) for 24 hours (Figure 6c-d). These findings suggest that Ndufs4 KO cells have a lower mitochondrial membrane potential and higher levels of cellular ROS than WT counterparts (Figure 6c-d), and that pre-treatment with NR mitigates these effects. Paced at 0.5 Hz, we measured Ca^2+^ transients in these iPS-CMs and found that Ndufs4 KO cells had a substantially slower decay rate (longer T50, time to 50% from peak) when compared with isogenic iPS-CM control (p<0.001) and this was significantly improved by NR (p<0.001, Figure 6e [right panel] and 6F). The amplitude of Ca^2+^ transients was not significantly different among all three groups (Fig 6e [left panel] and 6f). These findings suggest slower SERCA2 Ca^2+^ reuptake in Ndufs4 KO iPS-CM, that is significantly ameliorated by NR treatment. To confirm the findings in HEK293 cells, we performed patch-clamp to measure the inward depolarizing Na^+^ current (I_Na_) in Ndufs4 KO iPS-CMs and its isogenic control. As shown by the current-voltage curves (Figure 6g), I_Na_ density at -15 mV (Figure 6h) and area under the curve (AUC) of I_Na_ -voltage curves (Figure 6i), Ndufs4 knock-out reduced I_Na_ by ∼45% (p=0.088 for I_Na_ at -15mV and p=0.04 for AUC of current-voltage curves, Figure 6h-i). Pre-treatment with NR for 24 hours significantly enhanced I_Na_, especially in the Ndufs4 KO iPS-CMs (Figure 6g-i).

**Fig. 6.**
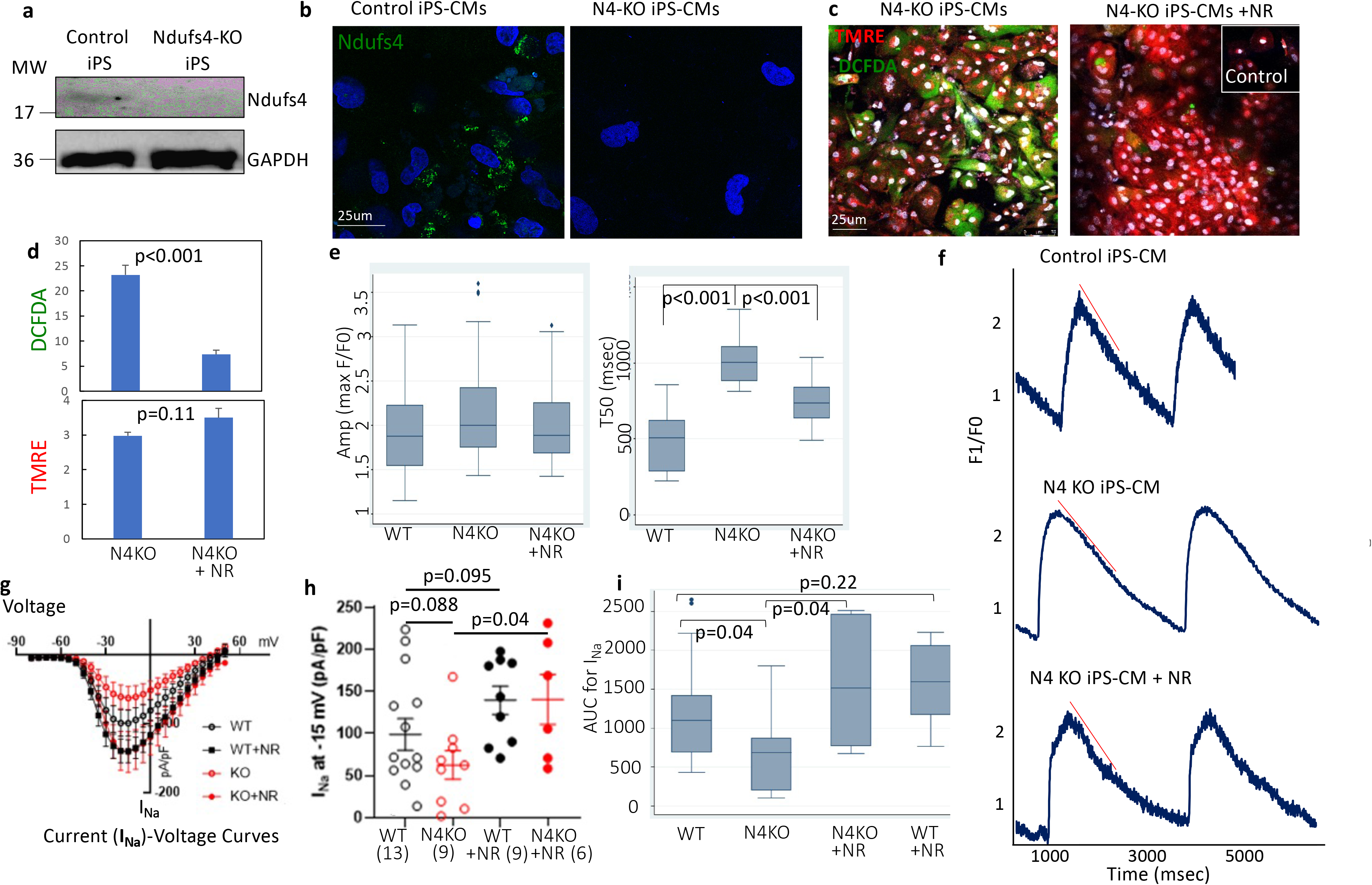
Deletion of Ndufs4 Decreased Na*_V_*1.5 current in human iPS-derived cardiomyocytes. (**a**) Representative immunoblots and (**b**) immunofluorescence images of WT and Ndufs4 KO human iPS-CMs immunolabeled for Ndufs4. (**c**) Live staining of control and Ndufs4 KO human iPS-CMs, with or without NR pretreatment, with TMRE (red) and DCFDA (green). (**d**) Quantitation of fluorescence in panel *c* (Data representative of three independent experiments). (**e-f**) Amplitude, decay rate (T50) and calcium transient in control and Ndufs4 KO human iPS-CMs with or without NR treatment, red line representing the slope

### Increased p53 acetylation and apoptosis in the Ndufs4-deficient brain

Our finding of enhanced apoptosis in the cerebellum and brainstem of Ndufs4^-/-^ mice and that this effect was ameliorated by NR supplementation (Figure 5j-u) suggested that protein hyperacetylation plays a critical role in apoptosis in the susceptible brain regions. Given that acetylation of p53 has been shown to enhance apoptosis (12), we examined iPS-derived neurons (iPS-N) for dysregulated NAD^+^/NADH and impaired Sirt1-dependent deacetylation of p53, which might potentially lead to neuronal apoptosis. Ndufs4 knock-out iPS and isogenic iPS controls were differentiated into neuronal progenitor cells (NPCs), which were then stimulated to form neural rosettes using an embryoid-body-based method (STEMCELL). The NPCs were allowed to mature into a mixed population of cells expressing a marker of GABAergic (the vesicular GABA transporter, VGAT in green, Figure 7a) or glutamatergic (vesicular glutamate transporter 1, VGluT1, in green, Figure 7b) neurons.

**Fig. 7.**
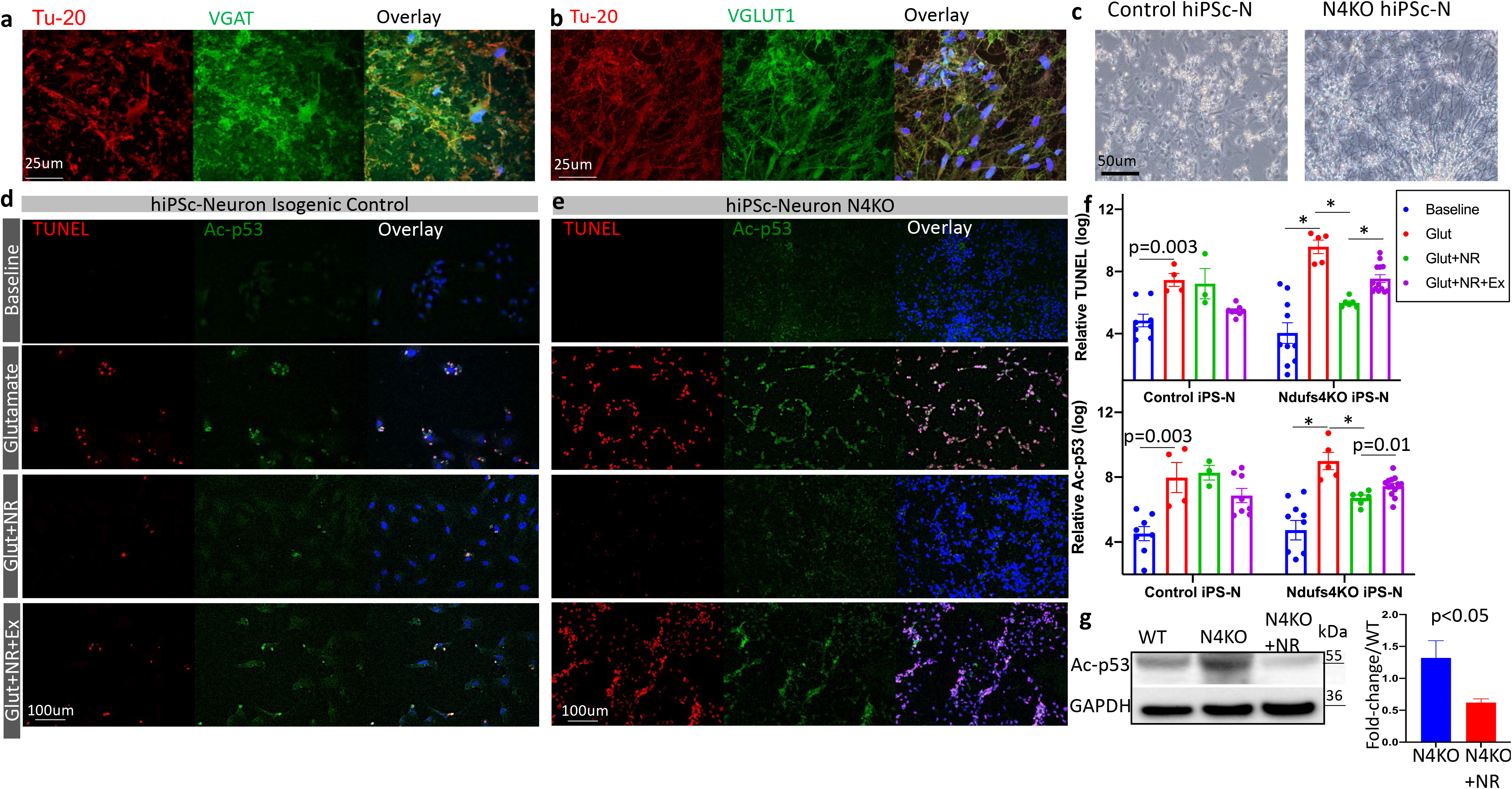
Human iPS-Neurons with Ndufs4KO are more susceptible to glutamate-induced apoptosis, mediated through acetylation of p53. (**a-b**) Characterization of human iPS-Ns by immunofluorescence for pan-neuronal marker β-Tubulin (Tu-20), vGAT (vesicular GABA transporter) or vGluT1 (vesicular glutamate transporter). (**c**) Morphology of mature human iPS-Neurons. (**d-e**) Representative fluorescence images of iPS-neurons treated with glutamate (30 μM), NR (1mM), and EX527 (10 μM). (**f**) Quantitation of relative fluorescence intensity of indicated groups in *d-e* (log-transformed for data normalization). (**g**) Representative western blot of K120-p53 in mouse cerebellum/midbrain from specified treatment groups and quantification (Data are representative of three independent experiments) (* p<0.001).

These VGAT or VGluT1 markers colocalized with the pan-neuronal marker beta-3 tubulin, Tu-20 (Figure 7a-b red), indicating a mixed population of iPS-Ns. At ∼30 days post-differentiation from neural rosettes (Figure 7c), these mixed iPS-Ns were stimulated with glutamate (30 µM), with or without NR (1 mM). The effect of glutamatergic excitotoxicity after 24 hours was assessed by TUNEL assay and measurement of acetyl-p53 (Figure 7d-e). Acetylation of p53 at either lysine K120 or K382 promotes p53-dependent apoptosis^12,13^ in response to DNA damage, cell stress, and oncogenic stress (14). Glutamate excitotoxicity led to a significant induction of apoptosis (TUNEL staining) in control iPS-Ns, in parallel with an increase in p53 acetylation (p=0.003). NR treatment with or without Ex-527, an inhibitor of Sirt1, in control iPS-Ns stimulated with glutamate did not significantly alter TUNEL or acetylated p53 staining (Figure 7d, f). In contrast, the glutamate excitotoxicity was potentiated in Ndufs4 KO iPS-Ns, shown by substantial increases in both TUNEL and Ac-p53 in response to glutamate stimulation (p<0.001, Figure 7e-f). Treatment with NR significantly attenuated the potentiation of p53 acetylation and TUNEL positivity in response to glutamate toxicity in the Ndufs4 KO iPS-N cells. Simultaneous treatment with Ex-527 and NR significantly attenuated the protective effect of NR on glutamate toxicity in Ndufs4 KO iPS-N cells. The partial inhibition of NR beneficial effect by Ex-527 suggests that the anti-apoptotic effect is mediated, at least in part, by NR-dependent Sirt1 deacetylation of p53. To confirm these findings *in vivo*, we performed Western blots of K120 acetylated-p53 in cerebellar/midbrain lysates. Acetylated p53 at K382 and K120 residues in cerebellar lysates from Ndufs4^-/-^ mice were increased compared with those from WT mice (Figure 7g, Supplemental Figure 8a-c). NR supplementation significantly reduced the hyperacetylation of p53, concomitant with the amelioration of neuronal apoptosis (Figure 5j-q).

## DISCUSSION

This study demonstrates novel mechanisms underlying cardio-encephalomyopathy in LS. Our analysis of mouse and iPS-CM models of this disease revealed that hyperacetylation of Na_V_1.5 promotes bradyarrhythmia and hyperacetylation of SERCA2a contributes to diastolic dysfunction. We further showed that the reversal of hyperacetylation by NR supplementation ameliorates these cardiomyocyte abnormalities in the context of Ndufs4 deficiency, *in vivo* and *in vitro*. Our study also provides a novel mechanistic explanation for the loss of neurons in LS, with involvement of p53 hyperacetylation and increased apoptosis in cerebellar and midbrain regions. Using targeted metabolomics, we demonstrated metabolic alterations in LS hearts and cerebellum/brainstem, characterized by significant increase in many amino acids and their metabolic derivatives, arachidonate and accumulation of TCA cycle intermediates (citrate, aconitate and isocitrate), decreased glutathione, niacin, and NAD^+^. We further showed that NR restores intracellular NAD^+^, glutathione and many of these metabolic changes. NR also attenuated p53 acetylation in iPS-Ns and the cerebellum and midbrain of LS mice, leading to an attenuation of neuronal apoptosis and mitigation of neuropathological lesions in LS brain.

Previous studies using germline LS mice focused largely on encephalomyopathy phenotypes(2). However, in clinical settings, ∼18-21% of LS patients have cardiac pathology, including hypertrophic cardiomyopathy and/or conduction abnormalities(3)(4). Thus, it is important to understand the cardiac manifestations in LS mice, which are not well characterized. Previous studies have utilized various muscle-specific loss of Ndufs4 to elucidate the role of mitochondrial complex I in cardiac physiology and metabolism. Although myocyte-specific loss of Ndufs4 (driven by CKM-NLS-Cre)(5) had been shown to lead to hypertrophic cardiomyopathy, cardiomyocyte-specific loss of Ndufs4 (driven by αMHC) did not show significant structural abnormalities or systolic dysfunction. Interestingly, in the latter group, pressure-overload induced heart failure was found to be aggravated(6) and associated with increased protein hyperacetylation due to alterations in the redox state and inhibition of Sirt3 activity(6). However, a more recent study of these mice (from the same group) showed that *ex vivo* ischemia-reperfusion injury was ameliorated in these mice, involved with a reduction of ROS, suggesting that ROS arising during reperfusion injury is generated mainly by complex I (15).

This study elucidates novel mechanisms that contribute to several phenotypes of cardio-encephalomyopathy in LS mice. First, we reported a novel finding of severe bradyarrhythmia, including SAN dysfunction (sick sinus syndrome) and occasional heart block that were associated with hyperacetylation of the K1479 in Na_V_1.5. Na_V_1.5 is expressed in the ventricular myocardium (16) and in the periphery of the sinoatrial node in which it is involved in the propagation of the action potential (17). Deletion of Ndufs subunits in heterologous system (HEK293) and iPS-CMs impaired various NAD^+^-dependent sirtuin deacetylases(9). Our previous study showed that membrane trafficking of Na_V_1.5 is disrupted by hyperacetylation of the K1479 residue in Na_V_1.5 in Sirt1^-/-^ mice, and this leads to reduced I_Na_ and impaired electrical activity in the heart (9). This study is the first to link mitochondrial complex I deficiency and the mechanisms underlying bradyarrhythmia with reduced NAD^+^/NADH, which leads to hyperacetylation of K1479 in Na_V_1.5 and subsequent reduction of the I_Na_, resembling the conduction block seen in MHC-Sirt1^-/-^ mice. The fact that NR reversed hyperacetylation of Na_V_1.5 at K1479 and enhanced I_Na_ in concert with restoration of normal sinus rhythm suggests that the hyperacetylation of Na_V_1.5 leads to bradyarrhythmia in the context of Ndufs4 deletion in LS hearts (Figure 8).

**Fig. 8.**
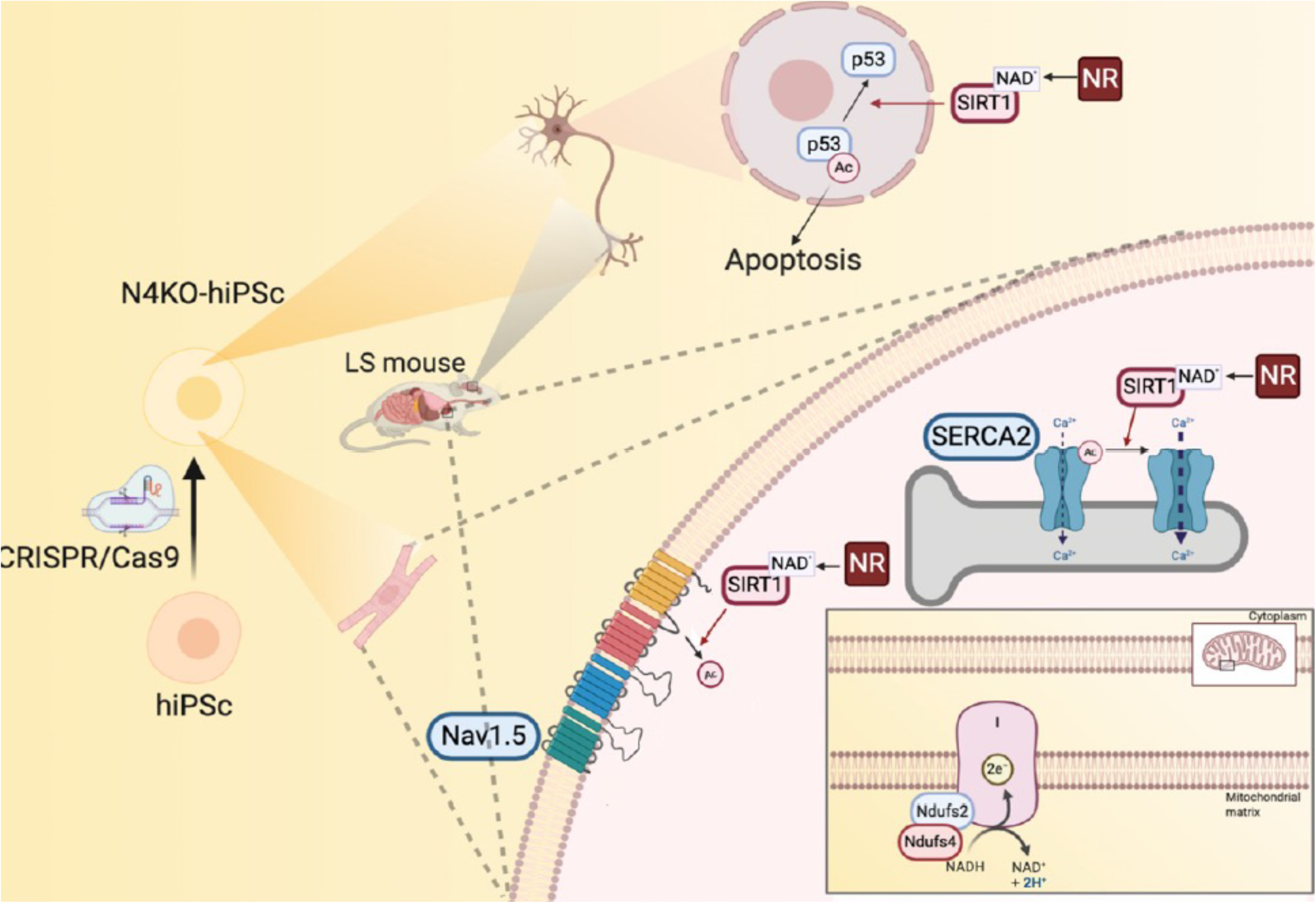
Schematic illustration of the molecular mechanism of cardio-encephalomyopathy in Ndufs4 KO mice and hiPSCs.

In addition to Sirt1-dependent Na_V_1.5 acetylation, our previous study showed an acetylation-independent increase in I_Na_ by NR supplementation through PKC phosphorylation of Na_V_1.5 (10)(18). Furthermore, a defect in mitochondrial complex I may also trigger oxidative and/or reductive stress (19), which has been shown to slow Na_V_1.5 inactivation, and/or to decrease the peak I_Na_, leading to arrhythmia (20)(21). Our mass spectrometric analysis in LS mouse hearts showed substantial reductions in reduced glutathione (GSH), a major reducing substrate in the endogenous antioxidant system, which was also restored by NR supplementation, suggesting that the beneficial effect of NR may also be mediated, at least in part, by restoring GSH (Figure 1c).

The second finding in our study that sheds light on LS-associated cardiomyopathy is that, despite the absence of hypertrophic cardiomyopathy or systolic dysfunction, LS mice exhibited significant diastolic dysfunction in association with hyperacetylation of SERCA2a. Hyperacetylation of SERCA2a was recently shown to reduce Ca^2+^ reuptake into SR (i.e., SERCA2a dysfunction) (22), predisposing to the development of heart failure. Increased SERCA2a acetylation was also reported in various failing heart samples, including end-stage human cardiac explants, murine hearts with pressure-overload induced heart failure, and porcine models of myocardial infarction. These observations suggest that increased acetylation of SERCA2a might be a shared mechanism leading to impaired SERCA2a function in failing hearts (22). Our finding that NR reversed the hyperacetylation of SERCA2a and ameliorated its compromised function in LS mouse hearts and Ndufs4-deficient iPS-CMs suggests that SERCA2a hyperacetylation may contribute to diastolic dysfunction in LS mice.

The third key finding in our study of LS-encephalomyopathy is the substantial loss of neurons, increased neuronal apoptosis and increased infiltration of the cerebellum and midbrain by activated microglia, concomitant with an increase in acetylated-p53. Acetylation of p53 has been shown to mediate activation of the downstream effector p21, leading to apoptosis (14). Our demonstration of the beneficial effects of NR in decreasing acetylated-p53 and neuronal apoptosis (both *in vitro* and *in vivo*), including attenuating microglial activation and overall neuropathology in the LS mouse brain, suggests that activation/hyperacetylation of the p53-mediated apoptotic pathway contributes to the severe encephalopathy observed in LS mice (Figure 8).

Mechanistically, Sirt1-mediated deacetylation of p53 inactivates the sequence-specific transcriptional activity of the protein and represses p53-mediated arrest of cell growth and apoptosis in response to DNA damage and oxidative stress. This process also prevented p53-dependent transactivation of p21 (23). Several lysines in human p53 (K120, K370, K372, K373, K381, K382) are acetylated following various forms of DNA damage, enhancing the transcriptional activity of p53 and regulating the fate of the cell (13). As we have shown, acetylation of p53 at K120, within its DNA-binding domain, is crucial for p53-mediated apoptosis. Our findings are reinforced by a previous study demonstrating that Sirt1 plays a neuroprotective role in the context of brain ischemia through deacetylation and subsequent inhibition of p53-induced and nuclear factor B-induced κ apoptotic and inflammatory pathways (24). Similarly, the Sirt1 mediated brain-protective effect of Maresin1 (macrophage mediator in resolving inflammation) in a mouse model subjected to middle cerebral artery occlusion demonstrates the involvement of Sirt1 signaling in the reduction of mitochondrial damage in the context of cerebral ischemia reperfusion injury (25).

The role of protein hyperacetylation in heart failure has been controversial. Increased global, mitochondrial, and metabolic protein acetylation has been documented in αMHC-Ndufs4^-/-^ mouse hearts, human heart failure and several experimental models, suggesting protein hyperacetylation is critical in the pathogenesis of heart failure. In contrast, a recent study showed that extreme acetylation of the cardiac mitochondrial proteome in carnitine acetyltransferase and Sirt3 double knock-out (DKO) mice did not potentiate pressure-overload induced heart failure. Even though the mitochondrial acetyl-lysine landscape of these DKO hearts was elevated to levels much higher than those observed in response to pressure overload or Sirt3 KO alone, deep phenotyping of mitochondrial function in the DKO revealed a surprisingly normal bioenergetics profile (26). The current study provides a direct mechanistic link between mitochondrial dysfunction and cardiomyopathic phenotypes caused by impairment of Sirt1 due to NAD^+^ deficiency. Our studies of LS mice elucidate a mechanism that underlies bradyarrhythmia and diastolic dysfunction which is promoted by hyperacetylation of Na_V_1.5 and SERCA2a, respectively. Both Na_V_1.5 and SERCA2a are critical cardiac proteins and well-known targets of Sirt1, rather than the targets of mitochondrial Sirt3. Collectively, our findings and previous publications suggest that it is the protein targets of acetylation themselves, rather than the degree of global protein acetylation, that play a critical role in cardiomyopathic phenotypes.

The fourth key finding is significant increase in many amino acids and certain TCA intermediates (citrate, aconitate, and isocitrate) in affected LS tissues, including cerebellum/brainstem and heart by targeted metabolomics. These are likely due to the impairment of various NAD^+^-dependent dehydrogenases, such as isocitrate dehydrogenase, of which the impairment may contribute to the accumulation of isocitrate, aconitate, and citrate. Likewise, decreased NAD^+^/NADH in LS may impair NAD^+^-dependent BCKD, which is responsible for BCAA oxidation, leading to accumulation of BCAAs (Figure 1a). NR supplementation significantly increased the NAD^+^ levels, decreased NAD^+^/NADH, and restored the metabolic derangement in LS cerebellum/brainstem and heart. Interestingly, several amino acids, particularly BCAAs, are potent inducers of the mechanistic target of the rapamycin (mTOR) signaling pathway, which has been shown to be activated in the LS brain, in concert with increased amino acids in the brainstem (27). Rapamycin, a specific inhibitor of mTOR, robustly enhances survival and attenuates disease progression. For example, it delays the onset of neurological symptoms, reduces neuroinflammation, and prevents brain lesions in these LS mice (28). A recent study reported that rapamycin restored mitochondrial protein levels and reduced the abundance and activity of multiple PKC isoforms in whole brain lysates of LS mice (29). Indeed, in a small clinical study, the mitochondrial disease phenotypes of four kidney transplant patients with mitochondrial disease (MELAS) improved after the immunosuppressant was replaced with rapamycin to achieve mTOR inhibition (30). However, because rapamycin is an immunosuppressant, the application to treat mitochondrial diseases might be limited by its side effects

A few treatment options have been proposed to alleviate LS pathologies in the same mouse model used in this study. Apart from rapamycin, chronic hypoxia has been shown to improve survival, body weight, body temperature, behavior, neuropathology, disease biomarkers and brain NAD^+^ concentration (31) in LS mice. In these studies, chronic hypoxia was achieved by putting the LS mice in a hypoxic chamber (32) or inducing severe anemia by phlebotomy or inhaling low-dose carbon monoxide (33). Although these proof-of-concept approaches are interesting, the clinical translation of these therapeutics may be challenging. A recent study using nicotinamide mononucleotide, NMN, also showed that this agent extended the lifespan of Ndufs4^flox/flox^ Meox2-Cre mice by restoring the NAD^+^ redox balance and lowering the accumulation of hypoxia-inducible factor 1-alpha in the skeletal muscle of Ndufs4 KO mice (34). However, no apparent beneficial effects were observed in the brain. Our current study elucidates the mechanism underlying the NR-mediated improvement of encephalopathy in Ndufs4KO mice, i.e., deacetylation of p53, and thus inactivation of the p53-dependent apoptotic pathway, leading to attenuation of neuronal apoptosis and the reduction of microglial activation in the cerebellum and the brainstem region.

The current study is the first to apply human iPS-CMs and iPS-Ns to model LS cardio-encephalomyopathy to elucidate mechanisms responsible for arrhythmia, diastolic dysfunction, and neuronal apoptosis. Using Ndufs4 KO iPS-CMs, we show that I_Na_ and SERCA2a function is decreased in these cells, and that these effects can be prevented by NR treatment. In iPS-Ns, Ndufs4 deficiency led to hyperacetylation of p53, resulting in increased neuronal apoptosis. The application of iPS modeling to human diseases has great potential to achieve the translation potential of personalized medicine and test the patient-specific efficacy of novel therapeutics. Given that NR is a vitamin B3 supplement with excellent safety profiles (35), our study provides robust evidence to support the potential clinical use of NR in LS patients.

## METHODS

### Study design

This study was designed to identify the mechanisms underlying cardio-encephalomyopathy in LS mice (germline Ndufs4^-/-^ deletion). Our hypothesis was tested both *in vitro* and *in vivo* using mice and human iPS cells modeling LS. Mice included germline Ndufs4^-/-^ (LS mice) and conduction tissue-specific Ndufs4^-/-^ (driven by HCN4-CRE). Six to 12 mice in each group were used to achieve a power of 0.8, with significance defined as p<0.05. The sample size per experiment is included in each figure legend. 21-30-days-old Ndufs4^-/-^ mice of both genders were given daily intraperitoneal NR injection (500mg/Kg/day) for 3-4 weeks.

For the *in vitro* studies, we used human embryonic kidney 293 (HEK293) cells and human iPS cells generated from a healthy Caucasian male (ATCC-1026) followed by deletion of Ndufs4 using the CRISPR/Cas9 method.

### Animal models

Germline Ndufs4^-/-^ mice were obtained from the University of Washington (2). The HCN4-Ndufs4 mice were generated by crossing Ndufs4^flox/flox^ mice with Hcn4tm2.1 (cre/ERT2) Sev/J mice (provided by Dr. Ivan Moskowitz, University of Chicago, available at JAX, # 024283 | HCN4CreERT2). All mice were on the C57/BL6/J background and were fed the regular diet from Harlan Teklad.

### Cell culture

HEK293 cells were obtained from the American Type Culture Collection (ATCC, Manassas, VA) and were cultured in 10% FBS and 1% PenStrep-supplemented DMEM medium.

### Generation of Ndufs knock-out cells using the CRISPR/Cas9 method

HEK293 cells were cultured to 70-80% confluence in 6-cm dishes. Cells were transfected with 2.5 μg of NF2SgRNA or NF4SgRNA (Addgene) and 5 μl of Lipofectamine 2000 (Invitrogen). Forty-eight hours after transfection, the cells were treated with the appropriate drug and selection was carried out for ∼7 days. The cells were then dissociated to single cells. After cells were counted, they were diluted serially in DMEM medium + 10% FBS to a final concentration of 5 cells per one well of 96-well plates and expanded for 2-3 weeks. Visible colonies were picked and reseeded in new wells for monolayer growth. Positive candidates were validated by western blotting.

### Maintenance of human iPS cells and differentiation of cardiomyocytes and neurons

Human iPS cells were cultured on Matrigel-coated plates (Corning, Life Sciences) and fed with a mixture of SFM and mTeSR^+^ at a ratio of 75%:25%. For cardiomyocyte differentiation, human iPS cells were passaged using gentle cell dissociation reagent (STEMCELL, catalog #07174) and replated at a density of 3.5*10^5^ cells/well on 12-well plates. Using STEMdiff^TM^ Cardiomyocyte Differentiation and Maintenance kits (STEMCELL, catalog #05010 & #05020), after cells reached 95% confluency, differentiation was initiated by replacing mTeSR with STEMdiff^TM^ Cardiomyocyte Differentiation medium supplemented with Matrigel and STEMdiff^TM^ Cardiomyocyte Differentiation Supplement A. Subsequently, cells were refed every two days, using STEMdiff^TM^ Cardiomyocyte Differentiation Supplement B & C, respectively. At Day 8, the medium was changed to complete STEMdiff^TM^ Cardiomyocyte Maintenance Medium, and it was changed every other day for 7 days. Cells typically began beating spontaneously on approximately day 8 after differentiation. Cardiomyocytes were purified from day 10 of differentiation using lactate method(sodium L-lactate, Sigma-Aldrich, USA).

For the generation of neural progenitor cells (NPCs), embryonic body (EB) protocol using STEMdiff neural induction medium and SMADi (STEMdiff™ SMADi Neural Induction Kit, Catalog#05835) was implemented. Briefly, aggrewell^TM^800 plates were prepared for experiments by pre-treatment of the wells with anti-adherence rinsing solution. Subsequently, each well was seeded with 3*10^6^ cells of single-cell suspension. Uniform EBs were observed in the aggrewell^TM^800 plates on day 1, ∼75% of the medium was replaced with fresh media every day for four days. EBs were harvested and replated at day 5 and medium was fully changed each day for six days. The percentage of cells induced to a neuronal fate was calculated on Day 8 based on the number of EBs with more than 50% neural rosettes divided by total number of EBs, and it was found to be ∼90%. On Day 12 neural rosettes were selected and replated, and full medium changes were performed daily, using STEMdiff^TM^ Neural Induction medium plus SMADi for another five days and the NPCs were passaged on Day 18. Neurons were generated from human iPS-NPCs using the STEMdiff^TM^ Neuron Differentiation kit (STEMCELL, catalog #08500) and STEMdiff^TM^ Neuron Maturation kit (STEMCELL, catalog #08510). Briefly, for neuronal differentiation we used a poly-L-ornithine/laminin coating and complete STEMdiff^TM^ Neuron differentiation medium. Neuronal precursors were seeded at a density of 3*10^4^ cells/cm^2^ and neurons were allowed to mature for at least one week in STEMdiff^TM^ Neuron Maturation medium or Brainphys^TM^ Neuronal medium (STEMCELL, catalog #05790) supplemented with Neurocult^TM^ SM1 Neuronal supplement (STEMCELL, catalog #05711), N2 supplement-A (STEMCELL, catalog #07152), recombinant human brain derived neurotrophic factor, recombinant human glial-derived neurotrophic factor, dibutyryl cAMP, and ascorbic acid.

### Electrocardiography and transthoracic echocardiograms

ECG recordings were obtained from conscious mice for at least 30 minutes, using the INDUS Rodent Surgical Monitoring system without anesthesia. Echocardiograms to measure ventricular size, wall thickness, and ejection fraction were performed on mice using the Vevo 2700 VisualSonics System (Toronto, ON, Canada). Prior to echocardiography, mice were injected with the sedative midazolam (with 0.1 mg subcutaneous injection). Cardiac images were obtained using a 30-MHz linear array transducer. Images of the parasternal short and long axis were obtained at a frame rate of ∼180-250 Hz. All image analysis was performed offline using the Vevo 2100 analysis software (v.1.5). Endocardial and epicardial borders were traced on the short axis view during diastole and systole. LV length was measured from endocardial and epicardial borders to the LV outflow tract in diastole and systole. The biplane area–length method was then used to calculate LV mass and ejection fraction. Diastolic function was measured using tissue Doppler imaging of the mitral annulus.

### Patch-clamp

Na^+^ current measurements were conducted using the whole cell patch-clamp technique as previously described (9). For voltage clamp studies, command pulses were generated using an Axopatch 200B patch clamp amplifier (Molecular Devices, San Jose, CA) and currents were sampled at 20 KHz through an A/D converter (DigiData 1440, Molecular Devices, CA) and low pass filtered at 5 kHz. Electrode offset potentials were zero-adjusted before a Giga-seal was formed. Fast- and slow capacitance was compensated. ≈ 85% series resistance was also compensated, yielding a maximum voltage error of ∼1mV. To minimize the effects of I_Na_ run-down on the results of the experiments, we carefully monitored the time-dependent change of I_Na_.

Recordings were started after the current reached a steady state, normally within 5 to 8 minutes. To record Na^+^ currents from HEK293 cells, electrodes of 2-3 MΩ were filled with a pipette solution containing (in mmol/liter) NaF 10, CsF 110, CsCl 20, EGTA 10 and HEPES 10 (pH 7.35 with CsOH), and the bath solution contained NaCl 40, 103 NMDG, KCl 4.5, CaCl_2_ 1.5, MgCl_2_ 1, and HEPES 10 (pH 7.35 with CsOH). For recording from human iPS-CMs, the bath solution contained 50 NaCl, 110 CsCl, 1.8 CaCl_2_, 1 MgCl_2_, 10 HEPES, 10 glucose, 0.0001 CdCl_2_ (pH 7.4 with CsOH). Patch pipettes were filled with 10 NaCl, 135 CsCl, 2 CaCl_2_, 3 MgATP, 2 TEA-Cl, 5 EGTA and 10 HEPES (pH7.2 with CsOH). All experiments were performed at room temperature (20-22 °C). Current-voltage (I-V) relationships were generated by plotting the current density elicited by voltage steps from -80 mV to +50 mV, at 5 mV intervals, from holding potential of -120 mV. Cellular capacitance was measured, and I_Na_ was normalized to cellular capacitance.

### Measurement of Ca^2+^-transient using confocal microscopy

Dissection of the SAN was performed in Tyrode solution (36°C) under the dissecting microscope. The SAN was delineated by the crista terminalis, the orifice of superior vena cava and the interatrial septum. Spontaneous beating Ca^2+^ or field-stimulated steady-state Ca^2+^ transients were measured in non-patched SAN cells or human iPS-CMs at ∼35°C. These cells were loaded with Rhod-2 AM (AAT Bioquest, Inc, Catalog #21062) at room temperature for 30 minutes, followed by de-esterification in Tyrode’s solution (containing 1.8 mM Ca^2+^) for 15 minutes. Line-scan confocal images were acquired at a sampling rate of 1.93 ms per line using a 63×, 1.3 NA oil immersion objective mounted on a Zeiss LSM 510 confocal microscope (Carl Zeiss MicroImaging GmbH). Steady-state Ca^2+^ transients were achieved by a 30-second pacing at spontaneous beating for the SAN at 0.5 Hz for human iPS-CMs. SR Ca^2+^ content was determined after steady-state stimulation at 1 Hz by measuring the amplitude of the Ca^2+^ release induced by local delivery of 20 mM caffeine. All digital images were processed using IDL 8.0 (Research System Inc).

### Immunoprecipitation and immunoblotting

Immunoprecipitation was carried out by incubating 4 μl of the acetyl-lysine antibody (Cell Signaling Technology, 9814) with 1 mg of tissue homogenate overnight, followed by adding 40 μl of protein A dynabeads (ThermoFisher) for 1 hour. After washing, immunoprecipitates were boiled in SDS-PAGE gel loading buffer, along with 500 μg of whole-cell lysates subjected to SDS-PAGE, transferred to nitrocellulose membrane and probed with a 1:500 dilution of the specified primary antibody and a 1:5000 dilution of peroxidase-conjugated secondary antibody.

Heart and cerebellar tissues were homogenized in RIPA buffer containing protease inhibitor cocktail (Roche). After quantification with BCA, equal amounts of proteins were loaded onto an SDS-PAGE gel, followed by standard immunoblotting. Chemiluminescent signal was developed using SuperSignal West Femto Maximum Sensitivity substrate (ThermoFisher, 34095), and blots were imaged using a GelDoc 2000 Chemi Doc system. Band densities were quantified using Image J (NIH).

### Antibodies

Primary antibodies used were against Na_V_1.5 (Alomone, 493-511), SERCA2a, custom designed anti-acetyl-K1479 Na_V_1.5 (YenZym Antibodies), Ndufs2 (Invitrogen, PA5-22364), Ndufs4 (ABclonal, A13519), p53 (ABclonal, A0263), Acetyl-p53 (Lys382) (Invitrogen, 710294), Anti-p53 (acetyl K120) (Abcam, ab78316), cleaved caspase-3 (Cell Signaling Technology, Asp175), VGluT1(Sigma-Aldrich, ZRB2374) and VGAT (Sigma-Aldrich, AMAB91043).

### Live cell staining and immunostaining

The HEK293 and iPS-CMs were plated on glass-bottom dishes. For live staining, the culture medium in the dish was exchanged with prewarmed (37°C) culture medium containing DCFDA (5mM) and TMRE (25nM), incubated for ∼30 minutes, then counterstained with Hoechst 33342 in new medium. For immunostaining, HEK293, iPS-CM and iPS-N cells were fixed in 4% paraformaldehyde, blocked, and incubated in primary antibodies overnight. They were then probed with secondary antibody and counterstained with DAPI. Images were acquired using a Leica SP8 confocal microscope.

### Metabolic chamber measurements

For analysis of whole-animal energy expenditure, animals were placed in the Oxymax CLAMS (Comprehensive Lab Animal Monitoring System, Columbus Instruments, Columbus, OH, USA) instruments. These cages have an open circuit system that directly measures various parameters over a 72 hour period, such as heat production, food intake, and movement (36). The Oxymax system has an open-circuit indirect calorimeter for lab animal research, allowing the measurement of oxygen consumption (VO_2_), respiratory exchange ratio (RER), and activity levels of mice. VO_2_ is a measure of the volume of oxygen used to convert energy substrates into ATP. The RER is calculated as the ratio of carbon dioxide production (VCO_2_) to oxygen consumption and can be used to estimate the fuel source for energy production based on the difference in the number of oxygen molecules required for the oxidation of glucose versus fatty acids. An RER of 0.7 indicates that fatty acids are the primary substrates for oxidative metabolism, whereas an RER of 1.0 indicates that carbohydrates are the primary energy substrates. Activity was calculated by summing the X-axis movement counts associated with horizontal movement ^35^. All protocols were approved by the University of Iowa Animal Care and Use Committee (IACUC). Heat production was calculated using the equation derived from Lusk (∼1928) to estimate aerobic respiration (Heat = 1.232*VCO_2_+3.815*VO_2_). VCO_2_ is described as the rate of carbon dioxide produced by the mouse, and VO_2_ is described as the rate at which oxygen is consumed by the mouse.

### Metabolomics measurements

All three groups of mice (WT, Ndufs4^-/-^, Ndufs4^-/-^+NR; n=4 each group) were euthanized. The cerebellum/midbrain and heart were rapidly frozen in liquid nitrogen within one minute of removal from the mice.

### GC-MS method

For metabolite extraction, the samples were extracted in ice cold 2:2:1 methanol/acetonitrile/water which contained a mixture of 9 internal standards (d_4_-Citric Acid, ^13^C_5_-Glutamine, ^13^C_5_-Glutamic Acid, ^13^C_6_-Lysine, ^13^C_5_-Methionine, ^13^C_3_-Serine, d_4_-Succinic Acid, ^13^C_11_-Tryptophan, d_8_-Valine; Cambridge Isotope Laboratories) at a concentration of 1 ug/ml each. The ratio of extraction solvent to sample volume was 18:1. Tissue samples were lyophilized overnight prior to extraction. Tissues were homogenized using a ceramic bead mill homogenizer, after the addition of extraction buffer. The samples were then incubated at –20°C for 1 hour followed by a 10-minute centrifugation at maximum speed. Supernatants were transferred to fresh tubes. Pooled quality control (QC) samples were prepared by adding an equal volume of each sample to a fresh 1.5 ml microcentrifuge tube. Processing blanks were utilized by adding extraction solvent to microcentrifuge tubes. Samples, pooled QCs, and processing blanks were evaporated using a speed-vac. The resulting dried extracts were derivatized using methyoxyamine hydrochloride (MOX) and N,O-Bis(trimethylsilyl)trifluoroacetamide (TMS) [both purchased from Sigma]. Briefly, dried extracts were reconstituted in 30 μL of 11.4 mg/ml MOC in anhydrous pyridine (VWR), vortexed for 10 min, and heated for 1 hour at 60°C. Next, 20 μL TMS was added to each sample, and samples were vortexed for 1 minute before heating for 30 minutes at 60°C. The derivatized samples, blanks and pooled QCs were then immediately analyzed using GC/MS.

GC chromatographic separation was conducted on a Thermo Trace 1300 GC with a TraceGold TG-5SilMS column (0.25 um film thickness; 0.25mm ID; 30 m length). The injection volume of 1 μL was used for all samples, blanks, and QCs. The GC was operated in split mode with the following settings: 20:1 split ratio; split flow: 24 μL/min, purge flow: 5 mL/min, Carrier mode: Constant Flow, Carrier flow rate: 1.2 mL/min). The GC inlet temperature was 250°C. The GC oven temperature gradient was as follows: 80°C for 3 minutes, ramped at 20°C/minute to a maximum temperature of 280°C, which was held for 8 minutes. The injection syringe was washed 3 times with pyridine between each sample. Metabolites were detected using a Thermo ISQ single quadrupole mass spectrometer. The data was acquired from 3.90 to 21.00 minutes in EI mode (70eV) by single ion monitoring (SIM). Metabolite profiling data was analyzed using Tracefinder 4.1 utilizing standard verified peaks and retention times.

We used TraceFinder 4.1 to identify metabolites in extracted samples, blank, and QCs. We do this by comparing sample metabolite peaks against an in-house library of standards. The standard library was prepared by processing and analyzing authentic standards via the method described above. We created a database of retention times and three fragment ions for each metabolite standard: a target peak/ion and two confirming peaks/ions. When running biological samples, metabolites are identified that not only match with the known retention times of the authentic standard, but also with its target and confirming peaks. Tracefinder was also used for GC-MS peak integration to obtain peak areas for each metabolite. After TraceFinder analysis, instrument drift over time is corrected for using local regression analysis as described by Li et al ^36^. Pooled QC samples which were run in duplicate at the beginning and end of the GC-MS run were used for this purpose. The data are then normalized to an internal standard to control for extraction, derivatization, and/or loading effects.

### Sample preparation for LC-MS

For metabolite extraction, samples were extracted in ice cold 2:2:1 methanol/acetonitrile/water which contained a mixture of 9 internal standards (d4-Citric Acid, 13C5-Glutamine, 13C5-Glutamic Acid, 13C6-Lysine, 13C5-Methionine, 13C3-Serine, d4-Succinic Acid, 13C11-Tryptophan, d8-Valine; Cambridge Isotope Laboratories) at a concentration of 1 μg/ml each. The ratio of extraction solvent to sample volume was 18:1. Tissue samples were lyophilized overnight prior to extraction. Tissues were homogenized using a ceramic bead mill homogenizer, after the addition of extraction buffer. Samples were then incubated at –20°C for 1 hour followed by a 10 min centrifugation at maximum speed. 400ul of supernatants were transferred to fresh tubes. Pooled quality control (QC) samples were prepared by adding an equal volume of each sample to a fresh 1.5 ml microcentrifuge tube. Processing blanks were utilized by adding extraction solvent to microcentrifuge tubes. Samples, pooled QCs, and processing blanks were evaporated using a speed-vac. The resulting dried extracts were reconstituted in 40 μL of acetonitrile/ water (1:1, V/V), vortexed and samples, blanks and pooled QCs were then analyzed using LC-MS.

### LC-MS Analysis

2 µL of metabolite extracts were separated using a Millipore SeQuant ZIC-pHILIC (2.1 X 150 mm, 5 µm particle size) column with a ZIC-pHILIC guard column (20 x 2.1 mm) attached to a Thermo Vanquish Flex UHPLC. Mobile phase comprised Buffer A – 20 mM (NH_4_)2CO_3_, 0.1% NH_4_OH and Buffer B: acetonitrile. The chromatographic gradient was run at a flow rate of 0.150 mL/min as follows: 0–21 min-linear gradient from 80 to 20% Buffer B; 20-20.5 min-linear gradient from 20 to 80% Buffer B; and 20.5–28 min-hold at 80% Buffer B ^37^. Data was acquired using a Thermo Q Exactive mass spectrometer operated in full-scan, polarity-switching mode with a spray voltage set to 3.0 kV, the heated capillary held at 275 °C, and the HESI probe held at 350 °C. The sheath gas flow was set to 40 units, the auxiliary gas flow was set to 15 units, and the sweep gas flow was set to 1 unit. MS data acquisition was performed in a range of m/z 70–1,000, with the resolution set at 70,000, the AGC target at 10^e6^, and the maximum injection time at 200 ms. Acquired LC-MS data were processed by Thermo Scientific TraceFinder 4.1 software, and metabolites were identified based on the University of Iowa Metabolomics Core facility in-house, physical standard-generated library. NOREVA was used for signal drift correction ^36^. Data per sample were then normalized to an internal standard (13C5-Methionine) to control for extraction, derivatization, and/or loading sample effects.

### Statistics

All analyses and calculations were performed using GraphPad Prism (version 8.0; GraphPad Software, Inc., CA, USA). The results are expressed as the mean ± standard error of the mean or proportion. Differences between groups were evaluated using Student’s t test, Kruskal Wallis test and analysis of variance (ANOVA) with Tukey’s test for comparison between two and multiple groups, respectively. Survival curves were drawn using the Kaplan-Meier method. To compare survival curves between groups, the log-rank test was applied. A p-value less than 0.05 was considered statistically significant.

## Data availability

The datasets generated and/or analyzed during the current study are included in this published article.

## Study approval

All animal experiments were approved by the Institutional Animal Care and Use Committee (IACUC) at the University of Iowa.

## Author contributions

JYY, DN, DFD performed experiments, data analysis and manuscript writing; YC, BC performed experiments and assisted in data analysis; MH performed neuropathological analysis, KI, LSS, BL, EDA and CB supervised the study, provided reagents and provided critical revision and editing of the manuscript; DFD developed the concept and design of the study. The order of co-first authorship is assigned by alphabetical order of the authors’ name.

## Acknowledgments

We thank the Fraternal Order of Eagles Diabetes Research Center Metabolomics Core Facility in the Carver College of Medicine at the University of Iowa for assistance in the measurement of metabolites. This research is supported by National Institutes of Health (grant reference number K08 HL145138 to DFD, and R01 HL147545 to KI, CB, BL).

