## Supplementary files for "Metabolic rescue ameliorates mitochondrial encephalo-cardiomyopathy in murine and human iPSC models of Leigh syndrome"

### Supplemental Figures

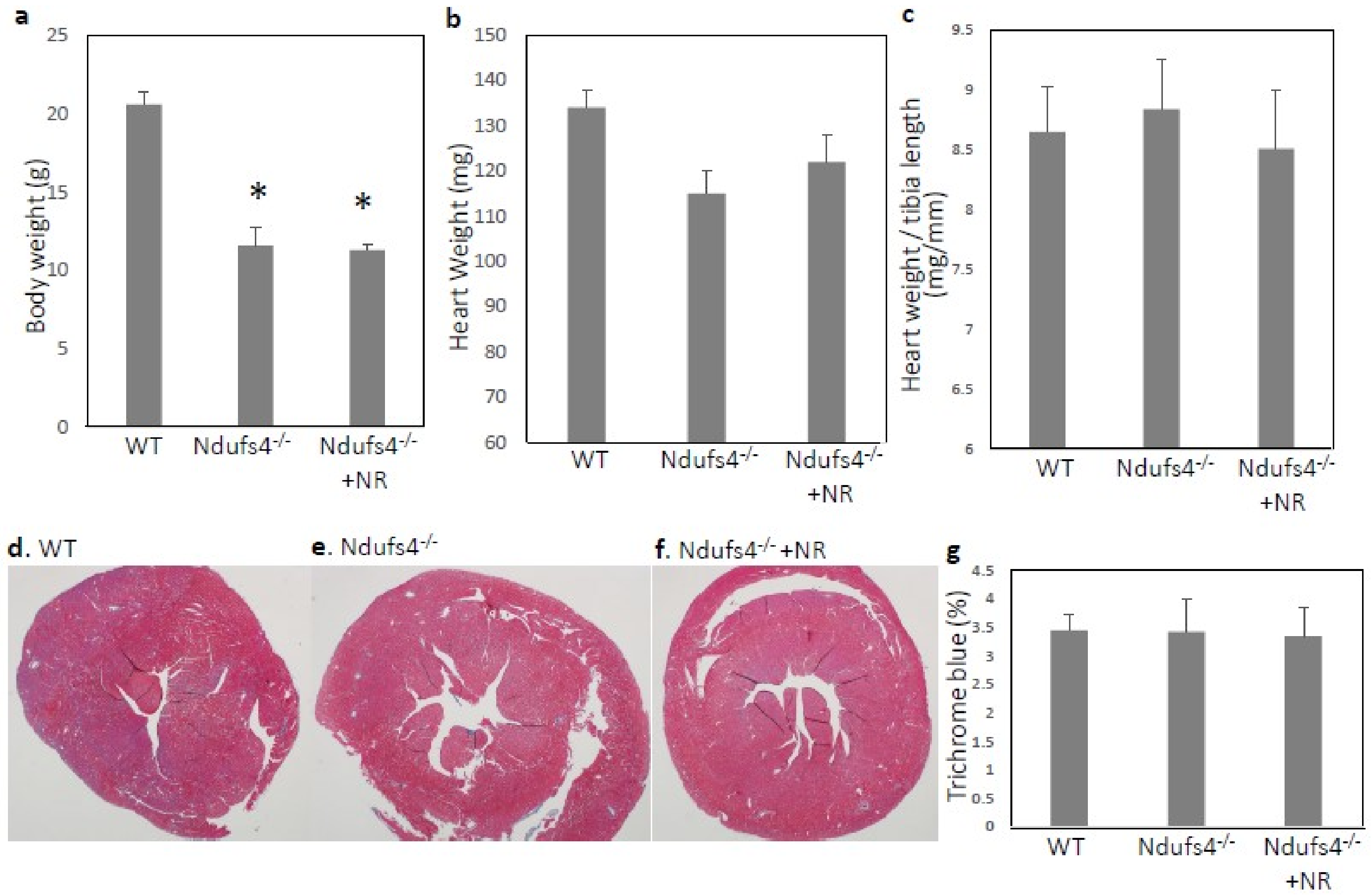


**Supplemental Figure 1. Body weight, heart weight, ventricle size and fibrosis of LS mice and WT littermate mice at approximately 6-7 weeks old.** (**a**) LS mice were runted with the average body weight ~60% of wild type (WT) littermates and post-weaning NR supplementation did not alter this. (**b**) Heart weight and (**c**) normalized heart weight was not significantly different from that in WT mice and this was not affected by NR. (**d-f**) Trichrome staining of left ventricles; (**g**) Quantitative analysis of trichrome blue area; *$p<0.05$ compared with WT. Data are mean ± s.e.m. of biologically independent samples. Statistical significance was determined by Student's t-test; n=5-7

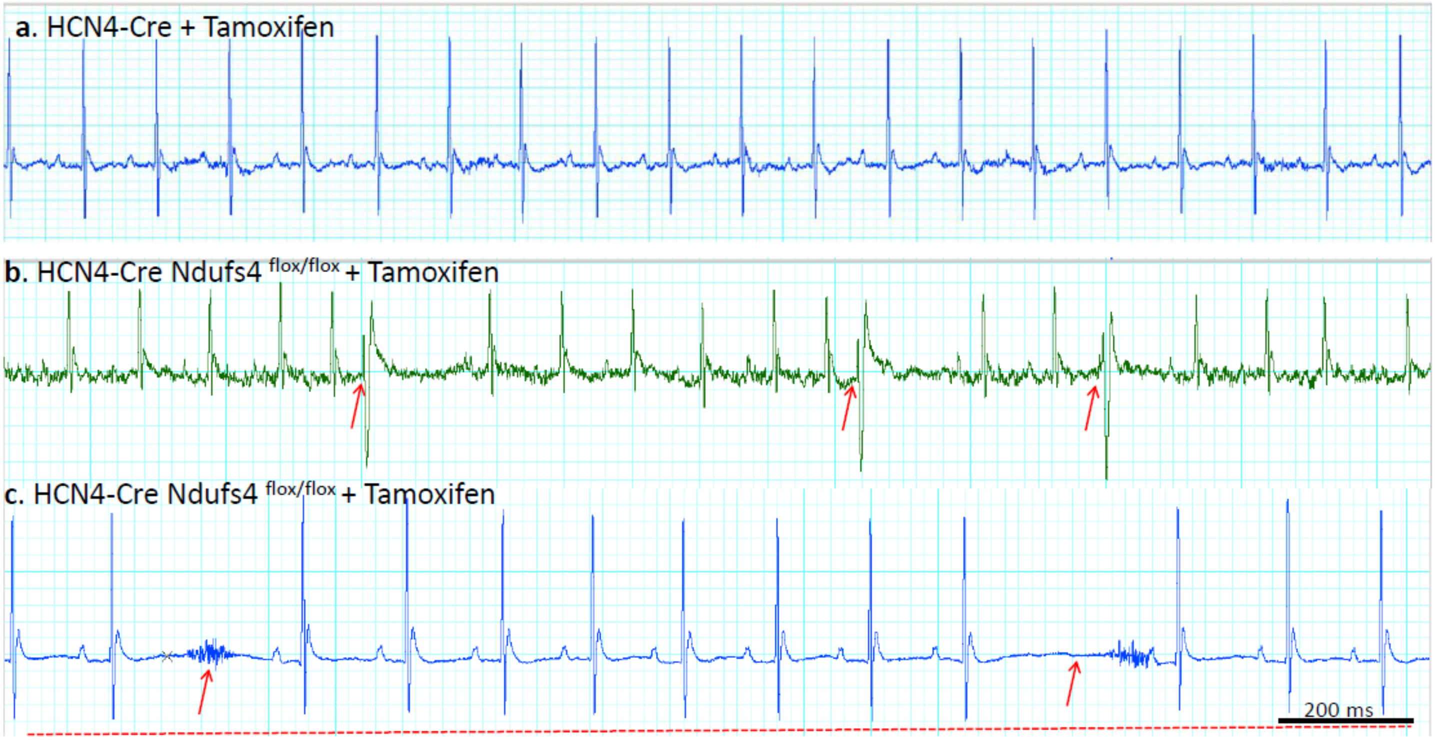


**Supplemental Figure 2. HCN4Cre/Ndufs4$^{-/-}$ mice developed various forms of cardiac arrhythmia**. (**a**) Normal sinus rhythm in HCN4-Cre injected with tamoxifen (n=3); (**b**) Premature contractions in 4 of 8 (50%) HCN-4-specific Ndufs4$^{-/-}$ mice, including APC, atrial arrhythmia and/or VPC; Examples of VPCs (red arrows) and APCs (blue arrows) are shown. (**c**) Sinus node dysfunction (arrows) and bradyarrhythmia (dotted line) was recorded in an additional 3 of 8 (37.5%) HCN4-Ndufs4$^{-/-}$ mice. One HCN4-Ndufs4$^{-/-}$ mouse (12.5%) showed normal sinus rhythm without arrhythmias.

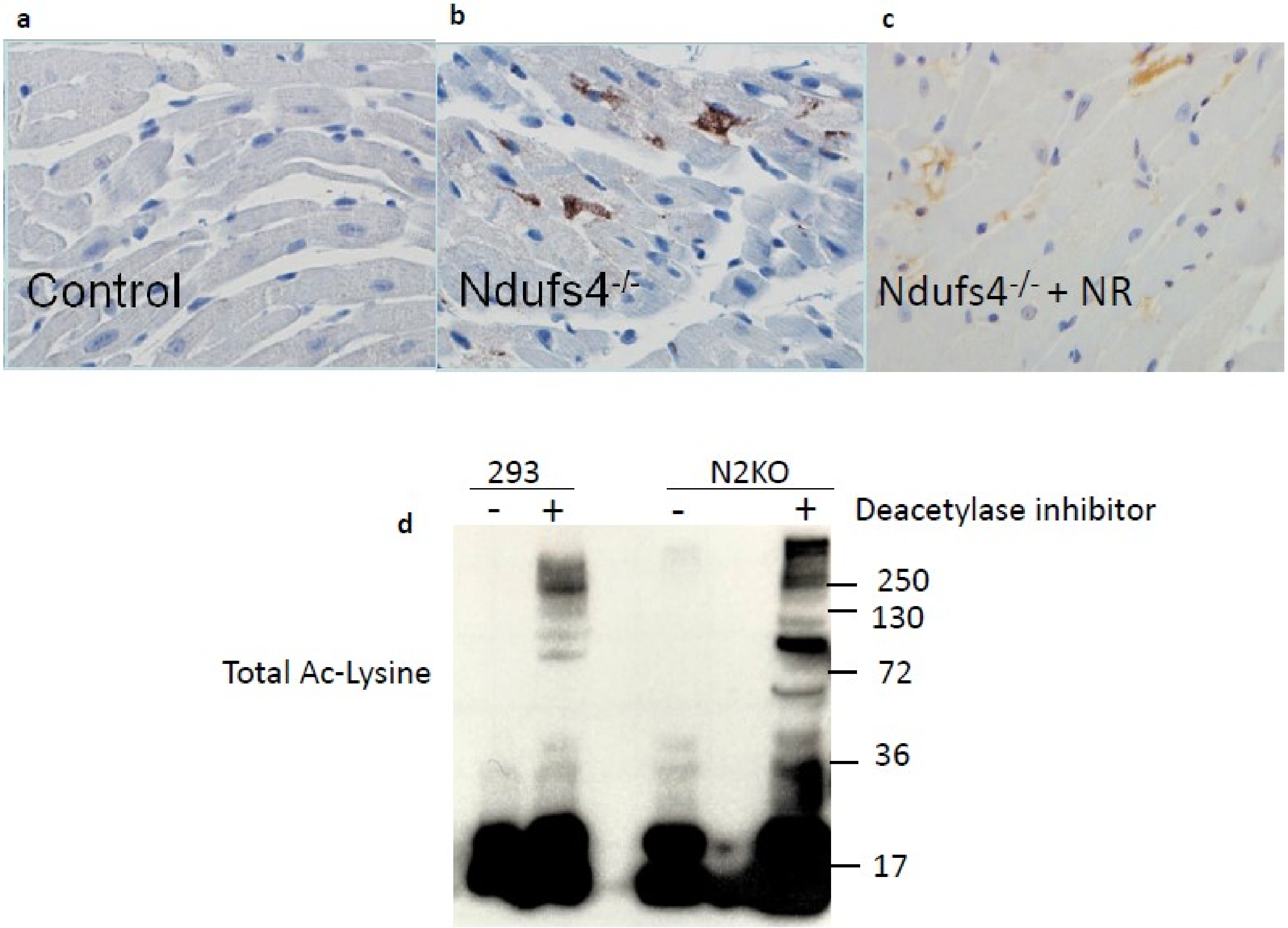


**Supplemental Figure 3. Increased acetylation of NaV1.5-K1479 in Ndufs4$^{-/-}$ heart.** (**a-c**) Immunohistochemistry staining of K1479 in NaV1.5 in left ventricular sections of WT and Ndufs4$^{-/-}$ mice before and after NR treatment. (**d**) Immunoblotting of total lysine acetylation in HEK293 and Ndufs2 knockout cells (n=3).

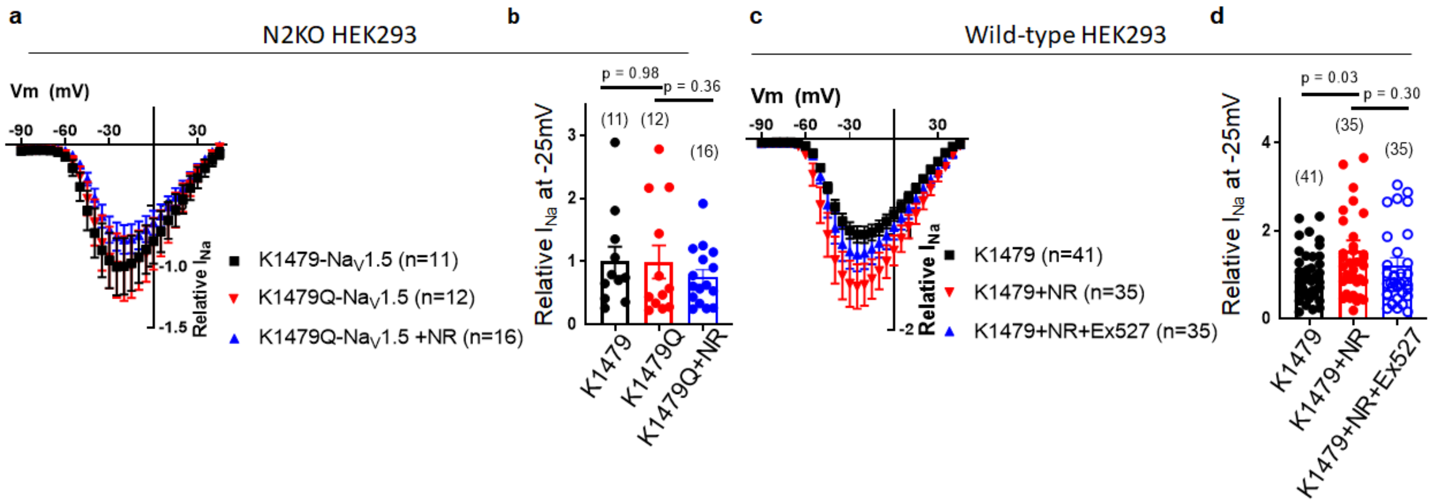


**Supplemental Figure 4. Sirt-1 mediated acetylation of K1479-Na$_V$1.5 is responsible for the effect of Ndufs2 KO or NR supplementation on $I_{Na}$. (a)** Normalized current–voltage relationships for Ndufs2 KO HEK293 cells expressing K1479-Na$_V$1.5 or mutant Na$_V$1.5 constructs mimicking acetylation (K1479Q). **(b)** Relative $I_{Na}$ density at –25 mV from data in panel *a*. **(c, d)** Normalized current-voltage relationships and relative $I_{Na}$ density at –25 mV for wild-type HEK293 cells expressing K1479-Na$_V$1.5 (control, vehicle) or treated with 5 mM NR with or without Sirt1 selective inhibitor Ex-527 (5 mM) for 48 hr. Data are mean ± s.e.m. of biologically independent samples. Statistical significance was determined by two-tailed Student's t-test.

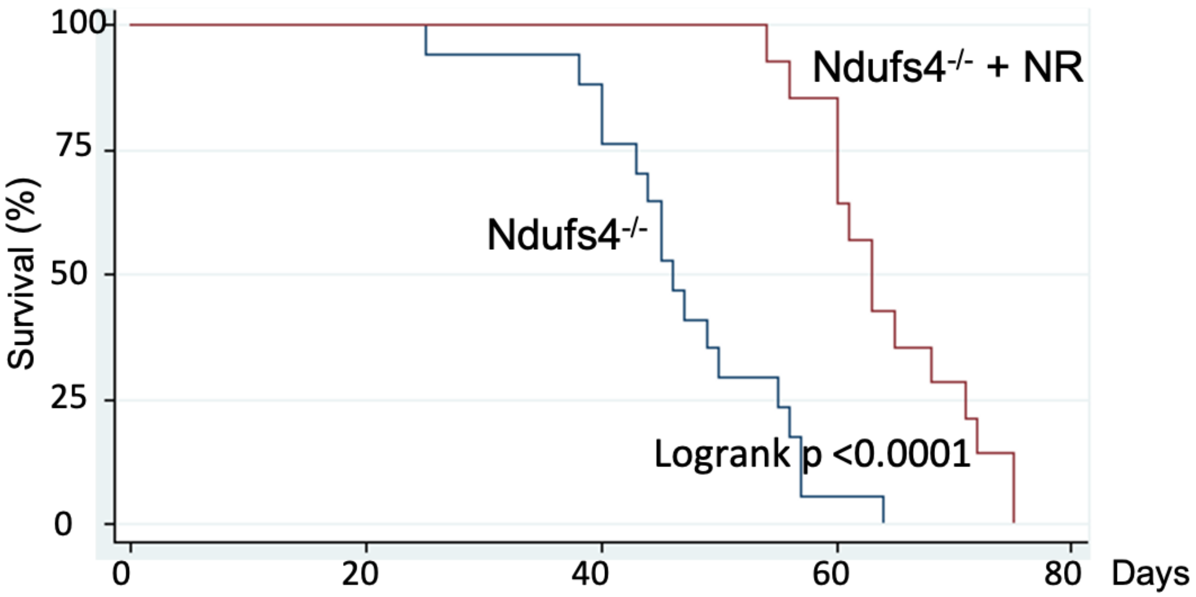


**Supplemental Figure 5. Survival curve of Ndufs4$^{-/-}$ mice**; n=14-17

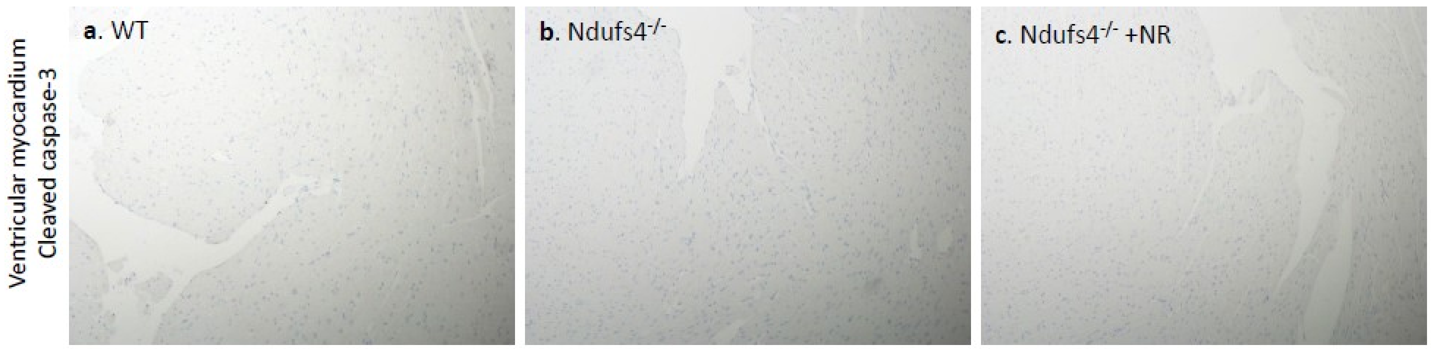


**Supplemental Figure 6. Immunohistochemistry of cleaved caspase-3 in mouse heart ventricles. Negative cleaved caspase-3 staining in all three groups**: (**a**) WT, (**b**) Ndufs4$^{-/-}$ and (**c**) Ndufs4$^{-/-}$ + NR; n=3.

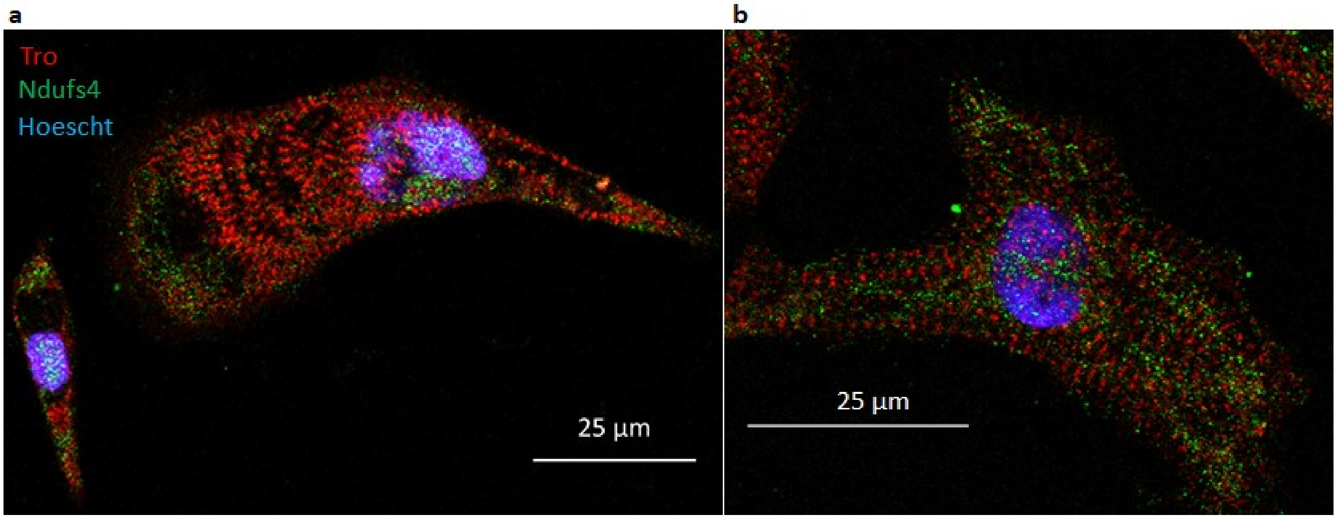


**Supplemental Figure 7. (a-b) Representative Immunofluorescence staining of iPS-CM** at ~Day 25 post-differentiation showing distribution of Ndufs4 and highlighting striated pattern of Troponin.

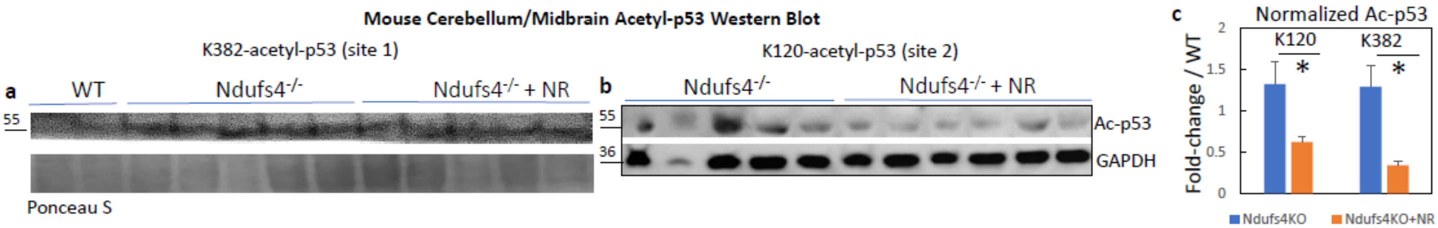


**Supplemental Figure 8. Acetyl-p53 in mouse cerebellum/midbrain from WT, Ndufs4KO and Ndufs4KO treated with NR.** (**a-b**) Representative western blot of K382-p53 and K120-p53 in mouse cerebellum/midbrain from specified treatment groups. (**c**) Quantification. (Data are representative of three independent experiments) (* $p<0.05$).

**Supplemental Movie legends**

**Supplemental Movie 1**. Ndufs4 KO untreated, showing marked decrease in activity, tremor, and cerebellar ataxia (rotational movement)

**Supplemental Movie 2**. Ndufs4 KO treated with NR, showing improvement of activity with the ability to walk straight with milder cerebellar symptoms (tremor and ataxia)

**Supplemental Movie 3**. Ndufs4 KO untreated, showing marked decrease in activity, tremor, and cerebellar ataxia (rotational movement)

**Supplemental Movie 4**. Ndufs4 KO treated with NR, showing improvement of activity with the ability to walk straight with milder cerebellar symptoms (tremor and ataxia)

**Supplemental Table legends**

**Supplemental Table 1**. Selective metabolites that increased or decreased by at least 20% in Leigh Syndrome hearts compared with wild type Hearts

**Supplemental Table 2**. Cardiac metabolites measured by GC/MS for WT, Leigh Syndrome (LS) and NR-treated LS mice (NR-LS)

**Supplemental Table 3**. Metabolites from cerebellum/brainstem measured by GC/MS for WT, Leigh Syndrome (LS) and NR-treated LS mice (NR-LS)
